# Phylogenetic Study of Long Non-Coding RNA *NEAT1* and *MALAT1* to Infer Structural-Functional Connection

**DOI:** 10.1101/2024.10.22.619594

**Authors:** Ksenia Arkhipova, Micha Drukker

## Abstract

*NEAT1* and *MALAT1* are the only two lncRNAs that utilise the tRNA-processing machinery for their maturation, coded very close to each other yet differing in function and evolutionary conservation. Here, we identified *NEAT1* and *MALAT1* orthologs in 545 mammals and studied their conservation patterns to gain functional insights. In the functional basis of *NEAT1Long*, we found G-quadruplexes and short sequence motifs that interact with the DSHS family of proteins and TDP-43, as well as gene length and self-complementary regions that further stabilise paraspeckle integrity. We found that transposable elements play a crucial role in *NEAT1* evolution by contributing structures recognised by DSHS proteins. We identified the *NEAT1Short* isoform in all *NEAT1* orthologs, and our results suggest its distinct functional role. Our findings also support the conservation of the isoform switch mechanism mediated by TDP-43. The function of *MALAT1* is likely based on its highly conserved primary sequence, and we demonstrated the gene regions under stronger purifying selection. This is the first large-scale phylogenetic study of *NEAT1* – a lncRNAs that lack sequence similarity between orthologs while maintaining functional and syntenic conservation.

## Introduction

Retrieving functionally important regions from an analysis of conservation patterns of primary and secondary structures of proteins and non-coding RNAs is a common approach. The method is based on the identification of conserved regions between orthologs, highlighting the pressure of purifying selection, which ensures that deleterious mutations are not established in the population, thereby maintaining only functionally essential structures (Charlesworth et al., 1993). Detailed mechanisms of function for two long non-coding RNAs, *NEAT1* and *MALAT1*, connected by the uniqueness of their maturation processes, are not yet clear. However, the association of these genes with neurodegenerative diseases and cancer highlights the urgent need to identify regions and properties crucial for their function.

*NEAT1* and *MALAT1* share unique structural elements at their 3’-ends and the maturation processing machinery. Both genes are located on chromosome 11 in the human genome, positioned in close proximity to each other and coded on the same strand (Fig. 1A), with SCYL1 adjacent to *MALAT1* and FRMD8 bordering *NEAT1*. Similar localisation of the genes in the mouse genome suggests synteny (Stadler, 2010). *NEAT1* is notably longer than *MALAT1*, spanning around 23 kilobases compared to *MALAT1*’s 8 kilobases. Similar to other lncRNAs, *NEAT1* and *MALAT1* are transcribed by polymerase II, but after that, unlike any other lncRNAs, tRNA-processing machinery is involved in their maturation. Specifically, the 3’-end of the genes forms a tRNA-like structure, which is recognised by RNase P and RNase Z, introducing two cuts before and after this structure, respectively (Fig. 1A) (Wilusz et al., 2008),(Sunwoo et al., 2009). After the tRNA-like structure is cut out, the newly formed 3’-end folds into a triple helix, which stabilises the gene (Brown et al., 2012). The tRNA-like structure (called mascRNA in the *MALAT1* gene) is further processed by another enzyme of tRNA maturation machinery - the CCA-adding enzyme, which can add two CCAs to the 3’-end of the structure instead of a single CCA, triggering its degradation. In five tested human cell lines it was demonstrated that *NEAT1*’s tRNA-like structures degrade in the cytoplasm, while mascRNAs remain stable (Wilusz et al., 2011), (Wilusz et al., 2008).

**Figure 1.**
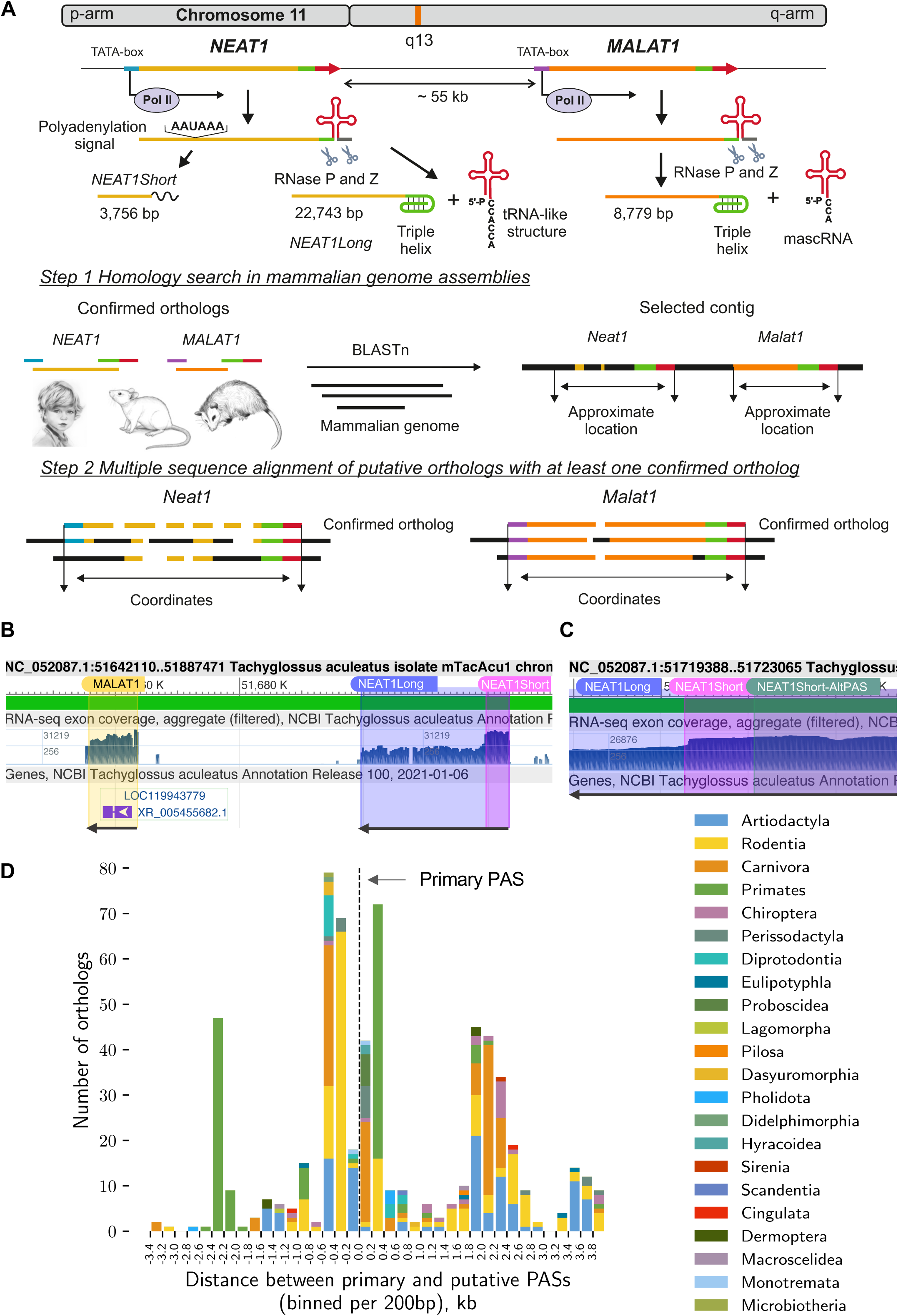
Identification of *NEAT1* and *MALAT1* orthologs. **A.** The organisation of *NEAT1* and *MALAT1* genes and the logic of ortholog coordinate identification. Promoter areas, TATA-boxes, tRNA-like structures, and triple helices are highlighted, with colours used uniformly throughout the scheme. Genomic regions lacking sequence similarity to confirmed orthologs are depicted in black. **B.** Confirmation of *NEAT1* and *MALAT1* ortholog predictions in *Tachyglossus aculeatus* (Monotremata order, short-beaked echidna)—the phylogenetically oldest species in our collection. Predicted coordinates were overlaid on mapped transcriptomic read profiles in the Genome Browser (http://genome.ucsc.edu). In the ‘Genes’ section of the Genome Browser, the automatically predicted genes identified in the region are shown. *Neat1* and *Malat1* are coded on the minus strand, with transcription direction indicated by arrows. **C.** Two predicted PASs in *Tachyglossus aculeatus*. Zoom-in view on the transcription profiles of *Neat1* in *Tachyglossus aculeatus* near the 3’-end of the *Neat1*Short isoform. The coordinates of the main and alternative PASs are overlaid. The primary PAS corresponds more closely to the drop in transcriptomic reads. **D.** Location and taxonomic distribution of alternative PASs in mammals. Most species possess an alternative PAS within 600 bp up- or downstream of the main PAS.

*NEAT1* has two isoforms, the long and the short, which we refer to as *NEAT1Long* and *NEAT1Short*, also known as *NEAT1*_1 and *NEAT1*_2, respectively. These isoforms share the 5’- end of the *NEAT1* gene, with the *NEAT1Short* forming a poly-A tail at approximately 3.3 kb of the gene. Several molecular factors regulating isoform balance have been identified. Among these, two proteins—HNRNPK and TDP-43—are residents of paraspeckles, which are formed by *NEAT1Long*. The accumulation of these proteins disrupts polyadenylation, triggering a readthrough of the gene and promoting the formation of *NEAT1Long* (Modic et al., 2019; Naganuma et al., 2012). In contrast, Integrator, a component of the polyadenylation machinery, enhances the production of the *NEAT1Short* isoform when recruited (Barra et al., n.d.). Although these mechanisms provide insights into isoform switching, it remains an open question whether *NEAT1Short* exists in all mammalian species encoding *NEAT1*.

The long isoform of *NEAT1* is an architectural nuclear-retained RNA, which is an essential component of paraspeckles (Sasaki et al., 2009). These nuclear bodies are dynamic ribonucleoprotein aggregates, built around *NEAT1* and stabilised by proteins of two main classes. Members of the DBHS family (NONO, SFPQ, PSPC1) are multidomain oligomerising proteins capable of binding nucleic acids (reviewed in (Knott et al., 2016)). They have a higher affinity for ssRNA than for dsRNA or ssDNA, while with less clear preferences for primary sequence patterns. NONO and SFPQ can also recognise secondary structures like stem loops, which can be formed from splice sites or inverted repeats of transposable Alu elements (IRAlu) or G-quadruplexes - guanine tracks separated by loops organised in layers by Hoogsteen hydrogen bonds ((Knott et al., 2016), (Arun et al., 2020), (Mou et al., 2022)). These proteins can form dimers with each other, enriching the diversity of interactions within paraspeckles. Another group of proteins that stabilize paraspeckles contain prion-like domains—low-complexity, unstructured regions, enriched in uncharged polar residues and glycine (Hennig et al., 2015). FUS and RBM14 are examples of essential paraspeckle proteins of this type (Hennig et al., 2015),(Fox et al., 2018). The conservation and importance of the individual elements of *NEAT1* recognised by these proteins remain unclear. It is also uncertain how interchangeable these proteins are, as not all proteins in these families have been identified as essential, with many considered merely important (Fox et al., 2018).

The formation of paraspeckles is linked to the transcription of *NEAT1* molecules. Initially, these molecules form aggregates, which then recruit multidomain proteins and other paraspeckle components (Mao et al., 2011). The paraspeckle structure consists of two main parts: the inner ‘core’ and the outer ‘shell’ (Hirose et al., 2019). These are distinguished by the folding of *NEAT1*, where the 3’ and 5’ ends are located in the ‘shell’, while the middle part of the gene forms the ‘core’, and by the predominant localisation of resident proteins (Hirose et al., 2019). While the paraspeckle structure is well established, the individual elements responsible for securing the distribution of resident proteins have yet to be identified.

Paraspeckles function by sequestering diverse proteins and RNAs, like IRAlu-containing mRNAs, miRNAs and transcriptional factors, which can affect gene expression (reviewed in (Wang et al., 2020; Yamazaki and Hirose, 2015)). Overall, an increase in paraspeckle numbers is associated with multiple neurodegenerative diseases (An et al., 2018), differentiation processes (Modic et al., 2019), and viral and bacterial infections. Paraspeckles are believed to be general organelles that appear in reaction to stressors (reviewed in (McCluggage and Fox, 2021)).

*MALAT1* is one of the most highly expressed genes in human cells. Like *NEAT1*, it is a nuclear-retained lncRNA. *MALAT1* is located in speckles – another type of nuclear body in direct contact with paraspeckles. Speckles contain factors involved in transcription regulation, mRNA maturation processes such as splicing and modifications, and mRNA export from the nucleus. *MALAT1* is a non-essential component of speckles (Nakagawa et al., 2012). Expression of *MALAT1* is associated with a range of cancers and diseases, as well as normal development (Song et al., 2021; Zhang et al., 2017),(Arun et al., 2020). Some of these functions are likely mediated by mascRNA, which may contribute to increased protein translation and proliferation by binding to a multi-tRNA synthetase complex (Lu et al., 2020).

It is widely accepted that transposable elements (TEs) play a significant role in mammalian evolution (Senft and Macfarlan, 2021). Several types of TEs are distinguished, among which endogenous retroviruses (ERVs), short interspersed nuclear elements (SINEs), and long interspersed nuclear elements (LINEs) are the most abundant in mammalian genomes (Osmanski et al., 2023). In primate genomes, Alu is the most abundant type of SINE (Deininger, 2011; Linker et al., 2017), capable of forming the aforementioned IRAlu structures, which trigger A-to-I modification. Intergenic lncRNAs are much more enriched in TEs compared to protein-coding genes (Hezroni et al., 2015). The most frequent TE type in lncRNAs is ERVs, while they are depleted in SINEs and LINEs (Kelley and Rinn, 2012). *NEAT1* is known to be enriched in repeats (Souquere et al., 2010), although their role in shaping *NEAT1* function and evolution has never been studied.

In general, lncRNAs exhibit poor conservation across vertebrates. It has been estimated that only 12% of human intergenic lncRNAs can be syntenically paired with orthologs from vertebrates, and around 21% of their transcript length can be aligned (Cabili et al., 2011). Another large study demonstrated that lncRNAs, on average, are more conserved at their 5’-ends. The study also identified 174 lncRNAs that were syntenic between many vertebrates but exhibited sequence similarity only among phylogenetically close organisms, raising questions about the function of these lncRNAs (Hezroni et al., 2015).

The information about the conservation of *NEAT1* is contradictory. In the early phylogenetic study on a diverse but concise set of around eight mammalian genomes, it was demonstrated that *NEAT1* orthologs are identifiable in Eutherians while absent in marsupials, likely due to incomplete assemblies (Stadler, 2010). However, between human and mouse *NEAT1* orthologs, there are only a few patches of similarity, although both form functional paraspeckles. Thus, *NEAT1* is an example of an lncRNA where a lack of sequence similarity does not imply a lack of function (Pang et al., 2006). This discrepancy became even more pronounced after the identification and functional confirmation of *Neat1* in opossum cells (marsupials), where traces of sequence similarity could be found in only 6% of the gene’s length. (Cornelis et al., 2016a). To date, a fourth mammal in which the *Neat1* gene has been identified and functionally confirmed is the naked mole-rat (Yamada et al., 2022), although, sequence homology of this ortholog was not analysed in detail. The conserved secondary structure could explain the ability of *NEAT1* to form paraspeckles. However, a comparison of the secondary structures of mouse and human short isoforms of *NEAT1* revealed predominantly different patterns, with only some regions of similarity. (Lin et al., 2018). Thus, the secondary structure of *NEAT1* likely does not contribute to paraspeckle formation and function. And the fundamental question of what elements are essential for *NEAT1* function remains unresolved.

Unlike *NEAT1*, *MALAT1* is a highly conserved lncRNA. Its orthologs were identified in zebrafish and other vertebrates (Stadler, 2010). Moreover, the conservation of a large part of the *MALAT1* secondary structure was demonstrated in 51 mammals (McCown et al., 2019). Additionally, the 3’-end unique structures (triple helix and tRNA-like) of *MALAT1* are very well conserved and exhibit co-evolving (simultaneous change of nucleotides in a complementary pair) patterns ((Stadler, 2010), (Monroy-Eklund et al., 2023).

In this study, we aimed to investigate the structure–function axis of *NEAT1* and *MALAT1* by identifying conserved regions, sequence features, structures, and regulatory elements. Since orthologs of these genes are not annotated in mammalian genomes, we began by developing a search algorithm that allowed us to locate the genes in 545 genomic assemblies of diverse mammals. We predicted *NEAT1Short* isoforms and alternative PASs, analysed the conservation of transcriptional regulation, triple helix elements, and tRNA-like structures. After analysing the overall similarity of *NEAT1* orthologs, we selected 16 the most dissimilar ones, which we called archetypes, for the identification of shared features. The primary sequence of the orthologs was scrutinized for nucleotide composition and the presence and enrichment of various repeats, like simple single-nucleotide repeats and transposable elements, and short sequence patterns. This analysis revealed the omnipresence of TG repeats and “TCTGTG” hexamers and high frequency of TEs integration. We also analysed secondary structure elements and identified G-quadruplexes as an essential component of all *NEAT1* orthologs, regardless of their primary sequence. Overall, our results suggest domains, elements, structures, and RNA processing of NEAT1 that are universally crucial for the function of paraspeckles.

## Results

### Defining genomic coordinates for *NEAT1* and *MALAT1* orthologs in 545 mammals

Since orthologs of *NEAT1* are annotated and confirmed in only three mammalian species (human, mouse, and opossum), and their primary sequences exhibit limited or no similarity, we undertook the challenge of identifying these orthologs in the available mammalian genomic assemblies. At the core of our algorithm lie the unique properties of this gene pair (Fig. 1A). Firstly, there is their co-localisation, as demonstrated in humans and mice. Secondly, although *MALAT1* orthologs are annotated only in human and mouse genomes, the gene is highly conserved. The initial homology search for *MALAT1* in mammalian assemblies, using the two known orthologs, supported the observation that the most of the gene can be easily located. We used these similarity patches of *MALAT1* as anchoring points for genomic contig selection, and the surrounding regions were then explored to locate *NEAT1*. By using the three *NEAT1* orthologs, we assumed that most species would exhibit some sequence similarity to at least one of the annotated orthologs. Due to the considerable length of the *NEAT1* gene, we separately searched for similarities to fragments of the promoter area containing the TATA-box and the triple helix followed by a tRNA-like structure. The high degree of conservation and individual alignment of the 3’-end structures constituted the third key component of our algorithm.

From the downloaded mammalian genomic collection, we reconstructed 506 *NEAT1* and 469 *MALAT1* gene orthologs, which were found evenly across all taxa without skew (Suppl. Fig. 1A). In a few genomes, we could not locate one of the two genes, either *NEAT1* or *MALAT1*, due to fragmented or incomplete genome assemblies. The identified *NEAT1* and *MALAT1* orthologs originate from 545 mammalian genomes (487 species, 122 families, 24 orders; Suppl. Fig. 1A), 17 of which belong to four orders of marsupials.

To substantiate our predictions, we analysed the transcription potential of genomic regions containing the identified orthologs by examining profiles of mapped transcriptomic reads in the Genome Browser ((Raney et al., 2024), Fig. 1B, C, Suppl. Fig. 1B). As a visual marker, we input the established coordinates of both genes (including the coordinates of the short isoform(s) of *NEAT1*) and compared our predictions with the results of transcriptome read mapping. We performed this verification for the most divergent *NEAT1* orthologs (archetypes), for which Genome Browser data were available, as these orthologs raise the most questions about the validity of our findings due to the lack of similarity in their primary sequences (Suppl. Fig. 1B). Since the remaining orthologs exhibit clear sequence homology to at least one of the archetypes, we assumed that the transcriptomic read mapping pattern would be comparable. In all tested cases, there was a very good agreement between our predictions and the expression profiles of the genomic regions for both genes.

To further support our predictions, we compared the identified orthologs of *NEAT1* and *MALAT1* in the naked mole-rat to recently published experimental evidence confirming their expression in most of the 14 tested tissues (Yamada et al., 2022). Although the gene sequences were not analysed in detail, we compared their starting coordinates with those identified in our study. We found that the coordinates mentioned in the publication are very close to our predictions, specifically, differing by 204 bp and 60 bp for *Neat1* and *Malat1*, respectively. This further highlights the validity of our programmatic approach.

One of the open questions in *NEAT1* biology is whether a short isoform is present and expressed in other mammals. Therefore, we attempted to predict *NEAT1Short* isoforms in the orthologs. Firstly, we identified the positions of the canonical polyadenylation signal (PAS), which comprises an ‘AATAAA’ motif. Next, we aligned the 3’-end of human *NEAT1Short* with other Eutherian orthologs in a multiple sequence alignment and searched for a predicted PAS within a narrow window of 220 bp. This method assumes a relatively high degree of sequence and length conservation of *NEAT1Short* among orthologs. Nevertheless, we successfully identified a single PAS in all Eutherians. We noted from the transcriptomic profiles (Fig. 1B, C) that *NEAT1Short* is distinguishable and forms a recognisable, twice-higher pattern. We also observed this in the phylogenetically oldest mammal in our collection, *Tachyglossus aculeatus* (order Monotremata, short-beaked echidna), where the predicted PAS corresponded well with the drop in transcriptomic read counts (Fig. 1C).

In the opossum (marsupial), Cornelis et al. found evidence for two active PASs approximately 500 bp apart (Cornelis et al., 2016a), and we used both in our prediction. With the exception of three genomes out of 17, we identified both sites in all marsupials and Monotremata. The predicted ‘main’ and alternative PAS were visualised in the Genome Browser (Fig. 1C). We found that the ‘main’ PAS is the more active one, as indicated by a sharper drop in transcriptomic reads. This is also consistent with the observations of Cornelis et al., where the two opossum sites exhibited different activity levels.

Next, we asked whether Eutherians also contain alternative PAS motifs in close proximity to the ‘main’ PAS or if this is specific to marsupials. We found that many other orthologs (n = 275, 55%) have alternative PASs in close proximity, located on both sides of the ‘main’ PAS (± 600 bp, Fig. 1D). The position of an alternative PAS is taxon-specific. Alternative PASs located 5’ to the main PAS were found in all marsupials and Monotremata, as mentioned above, in 72% of Rodentia orthologs, 37% of Carnivora, and 25% of Artiodactyla. The majority of orthologs with an alternative PAS downstream of the ‘main’ one belonged to 71% of Primates, 100% of Pholidota and Proboscidea, and 30% of Carnivora (Fig. 1D).

The assumption of *NEAT1* and *MALAT1* ortholog synteny was a key part of our search algorithm, and 92% of the 428 mammalian genomes analysed had both genes on the same contig. They were encoded in close proximity to each other, with the distance between them rarely exceeding 60 kb, averaging 36,755.3 ± 9,927.91 bp (Suppl. Fig. 1C). With the exception of two species (*Rousettus madagascariensis* and *Oryctolagus cuniculus*), both genes were encoded on the same strand of DNA. We suspect that the assembly quality of these two exceptions could explain this observation, as neither of these assemblies belonged to the GenBank reference set.

### Conservation of triple helix and tRNA-like structures of *NEAT1* and *MALAT1*

After identifying the orthologs of *NEAT1* and *MALAT1*, along with their isoforms and structural elements, we focused on comparing the 3’-end structures of each gene and assessing their similarity between *NEAT1* and *MALAT1* orthologs. These structures are unique to *NEAT1* and *MALAT1*, and while a high degree of sequence conservation has been demonstrated for structures of *MALAT1* orthologs, their variation in *NEAT1* remains unknown.

Overall, we found low divergence between the 3’-ends of *NEAT1* and *MALAT1* orthologs across all mammals (Fig. 2A). The triple helix structure consists of three principal parts: the structure-forming motif itself and two hairpin loops of different sizes (Fig. 2A). The conservation of the structure-forming motif is exceptional, as we found no mismatches in any orthologs of either *NEAT1* or *MALAT1*. However, the sequences of the hairpin loops varied and displayed clear specificity for *NEAT1* or *MALAT1*. In *NEAT1* orthologs, the sizes of both hairpin loops are almost the same (28.7 ± 0.98 bp, 29.8 ± 1.06 bp), but in *MALAT1* orthologs, the second hairpin loop is one-third shorter (31.36 ± 1.87 bp, 23.59 ± 1.8 bp, Fig. 2A).

**Figure 2.**
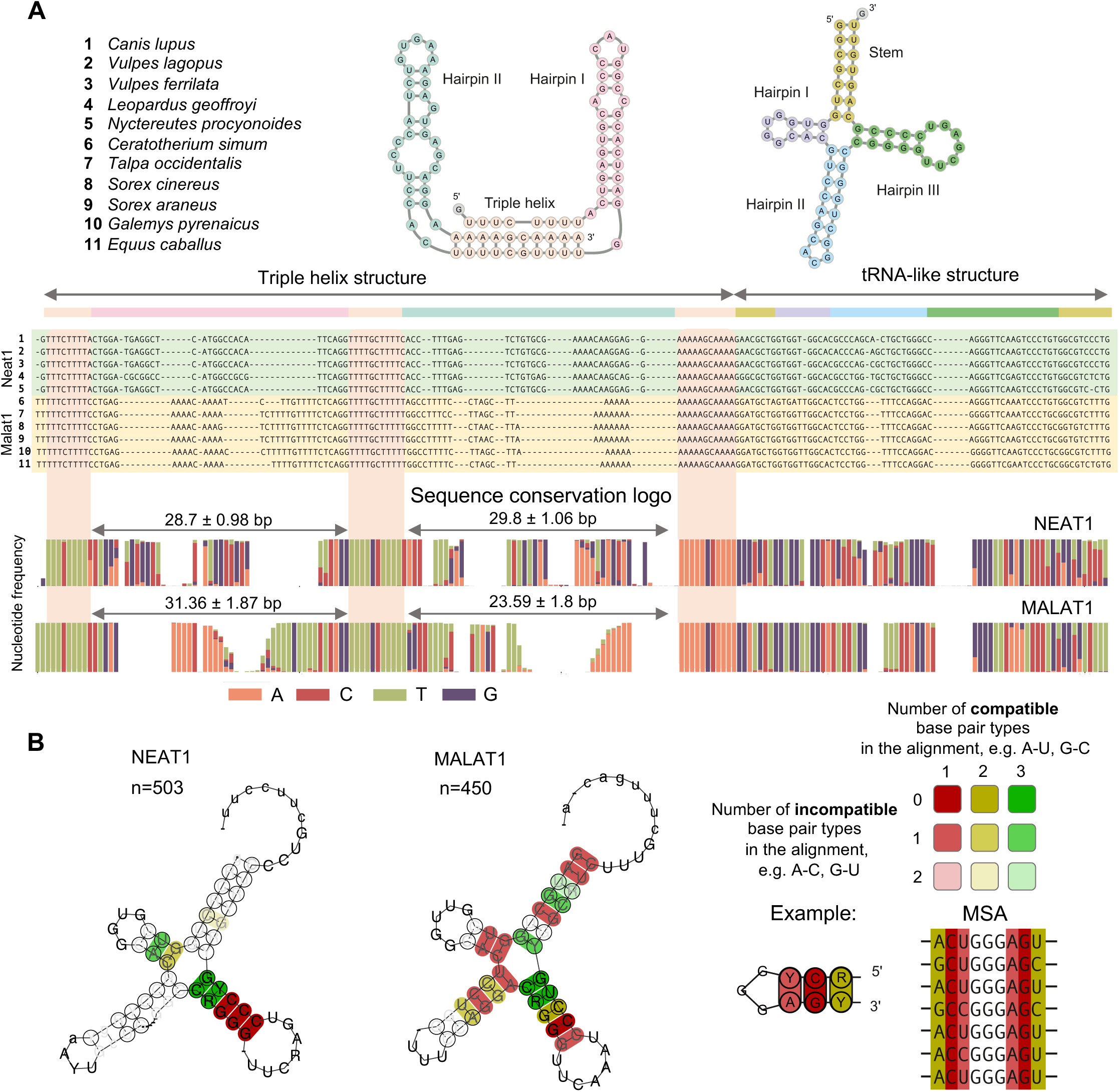
Conservation of 3’-end motifs of *NEAT1* and *MALAT1* orthologs. **A.** Secondary structure and sequence diversity of triple helices and tRNA-like structures in *NEAT1* and *MALAT1* orthologs. Secondary structures of the human triple helix and tRNA-like structure are shown at the top of the figure, with individual structural elements highlighted. Colours are used consistently throughout the figure. An example of the multiple sequence alignment of 3’-end structures of both *NEAT1* and *MALAT1* orthologs from the listed species is depicted. The summary of sequence diversity across all orthologs is presented as a coloured sequence conservation logo. The variance in length (mean ± std) of hairpins I and II of triple helices in *MALAT1* and *NEAT1* orthologs is specified. Highly conserved triple-helix-forming sequence regions are highlighted in both the alignment and logo figures. **B.** Co-evolving patterns of tRNA-like structures across all *NEAT1* and *MALAT1* orthologs in Eutherians. The most conserved base pairs are shown in dark red. High-intensity yellow and green indicate perfectly matching alternative base pairs (coevolving) in the MSA. The co-evolving patterns of the tRNA-like structure of *MALAT1* exhibit a much higher level of conservation in the whole secondary structure, while the tRNA-like structure of *NEAT1* mainly involves hairpin III with a highly variable hairpin II.

The degree of conservation in the primary sequence of the tRNA-like structures of *NEAT1* and *MALAT1* orthologs was high, although *NEAT1* orthologs exhibited slightly greater variation (Fig. 2A). We analysed patterns of coordinated nucleotide changes in complementary pairs (coevolving) to assess the pressure of purifying selection on the secondary structure of the tRNA-like elements. We found that the secondary structures of both genes are well conserved (Fig. 2B), and the sizes of individual elements, such as hairpin loops, do not vary drastically. Our analysis clearly highlighted the strongest purifying selection on the third hairpin loop of the tRNA-like structures in both genes, suggesting it has higher functional importance.

### Analysis of promoter and transcriptional control of *NEAT1* and *MALAT1*

The conservation of promoter regions and binding sites for transcription factors across different species can highlight the importance of a gene within certain gene networks and physiological processes. Conversely, variability in transcriptional regulation can suggest functional diversity of a gene in different taxa. We began with an analysis of the TATA-box and the downstream promoter area. Overall, this region was more conserved in *NEAT1* orthologs than in *MALAT1*, which is remarkable given the much higher primary sequence variability of *NEAT1* in mammals compared to *MALAT1* (Fig. 3A). We found that *NEAT1* orthologs in all Eutherians possessed the classical TATA-box sequence TATAAA, with greater promoter area diversity observed only in marsupials. Notably, *NEAT1* orthologs from the Chiroptera order exhibited an extra stretch of adenines downstream of the TATA-box. The variability of the transcription initiation site in *MALAT1* was significantly higher, and it was less variable only within individual mammalian taxa (e.g., Primates, Chiroptera in Fig. 3A). We also noted a higher diversity of TATA-box motifs in *MALAT1*, such as ‘CATAAA’ in the Chiroptera order, and both ‘AATAAA’ and the classical ‘TATAAA’ in Primates.

**Figure 3.**
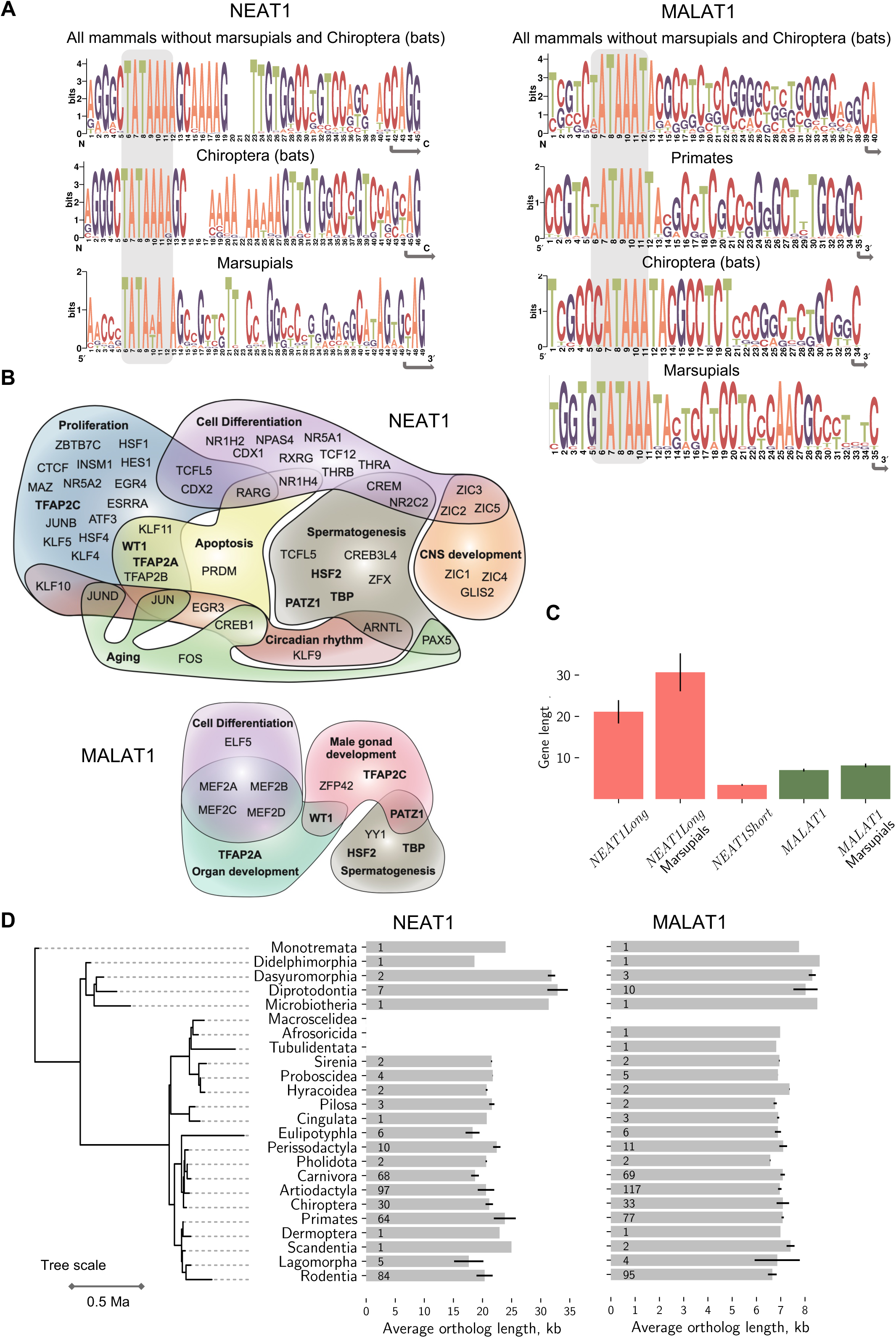
Transcriptional regulation of *NEAT1* and *MALAT1* orthologs and their length distribution. **A.** Conservation of TATA-boxes and promoter areas in *NEAT1* and *MALAT1* orthologs (sequence logo). TATA-boxes are highlighted with grey boxes, and the transcription start site is marked by an arrow. **B.** The most frequent GO terms associated with transcription factors, the binding sites of which were identified in at least 65% of orthologs of *NEAT1* and *MALAT1*. Shared biological processes associated with the same TF are depicted as overlaps. **C.** Average ortholog length and its variation across mammals. Marsupials exhibit the longest average length for *Neat1* and *Malat1*, while the *NEAT1Short* isoform shows much smaller length variation compared to *NEAT1Long*. Only non-gapped ortholog assemblies are taken into account. **D.** Length distribution of *NEAT1* and *MALAT1* orthologs in mammalian orders, arranged along a time-scaled phylogenetic tree. Only orthologs with non-gapped gene assemblies were used. The number of orthologs used for the assessment is indicated on the bars.

As a next step, we predicted transcription factor (TF) binding sites within the 1 kb promoter area of *NEAT1* and *MALAT1* orthologs, analysed the diversity of identified TFs, and explored the physiological processes in which *NEAT1* and *MALAT1* are potentially involved. An individual promoter of *NEAT1* and *MALAT1* orthologs had, on average, 216.4 ± 33 and 168.2 ± 29 TF binding sites, respectively. Although the average number of sites did not differ drastically, we investigated how many of these sites were identified across the majority of the studied orthologs. We observed that only a small number of TF binding sites were shared among the promoters of *MALAT1* orthologs. We applied a rather permissive threshold, defining ‘the majority’ as 65% of orthologs, resulting in 25 TFs for *MALAT1* and 123 TFs for *NEAT1* (Suppl. Table 1). This suggests a higher regulatory conservation of the *NEAT1* gene compared to *MALAT1*, which is noteworthy given the much higher primary sequence conservation of *MALAT1* orthologs.

Among the predicted TF binding sites for *NEAT1* and *MALAT1* orthologs, we identified 15 that overlapped: KLF16, PATZ1, PLAG1, SP4, TRAF2A and TRAF2C, WT1, ZFP14, ZNF281, ZNF454, ZNF460, ZNF701 and ZNF740, including EGR1 and SP1, which have been experimentally validated (Kumar and Mishra, 2022), (Binder et al., 2023), (Li et al., 2015), (Tian et al., 2023), (Che et al., 2021). Finally, we annotated the predicted TFs with GO terms, and a summary of the biological processes in which they are involved is presented in Fig. 3B. In addition to the well-established associations of *NEAT1* and *MALAT1* with certain pathways, our analysis suggests a novel role for this gene pair in spermatogenesis. This physiological process, along with cell differentiation, was shared between *NEAT1* and *MALAT1*. Overall, our results indicate a higher degree of conservation of the regulatory elements of *NEAT1* transcription compared to *MALAT1*.

### Gene length variation of *NEAT1* orthologs

*NEAT1* is the longest lncRNA in the human genome (Derrien et al., 2012), and its length may be a functionally relevant physical parameter for an architectural RNA, facilitating phase separation and stabilising paraspeckles. We analysed the distribution of lengths of *NEAT1* orthologs and found that the average length was 21,114.1 ± 2,811.3 bp (only assemblies without gaps were used), which is slightly shorter than the human *NEAT1* but longer than mouse *Neat1*, 22,743bp and 20,770bp, respectively. However, the difference between the longest and shortest variants was substantial: 14,505 bp in *Ochotona curzoniae* (plateau pika, Lagomorpha) and 36,456 bp in *Gymnobelideus leadbeateri* (Leadbeater’s possum, Diprotodontia). Notably, the lengths of *NEAT1Short* isoforms varied within a much narrower range, 3,415.18 ± 218.9 bp (Fig. 3C), which suggests potential functional importance.

We observed that the length of the *NEAT1Long* isoform and its variation exhibited some taxon-specific patterns (Fig. 3C, D). Marsupials from the Microbiotheria, Diprotodontia, and Dasyuromorphia orders had the longest *Neat1* genes of all mammals, averaging 30,659.9 ± 4,575.1 bp. However, we did not identify an association between gene length and taxon position in the phylogenetic tree (Spearman’s rho = −0.06, p = 0.18, correlation of *NEAT1* length vs phylogenetic distance from *Tachyglossus aculeatus*, Monotremata). Additionally, *NEAT1* length varied more within some orders, such as Primates and Artiodactyla, compared to Carnivora. Although the observed variation in *NEAT1* ortholog lengths was significant, *NEAT1* remains an exceptionally long gene in all mammals, underscoring the importance of this parameter for its function.

The length of *MALAT1* orthologs varied within a narrower range than that of *NEAT1*, 6,986.8 ± 326.78 bp (Fig. 3D), with a taxon-specific pattern. Marsupials, like *NEAT1* orthologs, had the longest *Malat1* gene (8,124.25 ± 449.08 bp), while rodents exhibited the shortest *Malat1* gene (6,653 ± 176.3 bp). These findings highlight the higher conservation of the *MALAT1* gene compared to *NEAT1*.

### *NEAT1* and *MALAT1* orthologs primary sequence diversity and *NEAT1* archetypes

To identify conserved sequence domains potentially crucial for the function of *NEAT1* and *MALAT1* and to select appropriate analytical tools, we assessed the overall similarity of primary sequences between pairs of orthologs. We generated a heatmap (Fig. 4A) depicting the average nucleotide identity (ANI) between ortholog pairs in an all-vs-all comparison, with mammals ordered according to the phylogenetic tree. It is evident that *NEAT1* orthologs formed several clusters of higher homology (red and white coloured groups) with a strong phylogenetic signal, as blocks of red clusters correspond to mammalian orders. However, the similarity level of *NEAT1* orthologs between clusters was very low, in some cases barely exceeding 20-30% ANI. From this, we concluded that conventional methods of primary sequence analysis, such as multiple sequence alignment, are not suitable due to the high sequence diversity. Notably, the majority of *NEAT1* orthologs, the ones from Primates, Chiroptera, Carnivora, Artiodactyla, and Rodentia excluding the Muridae and Cricetidae families, exhibited a high level of homology.

**Figure 4.**
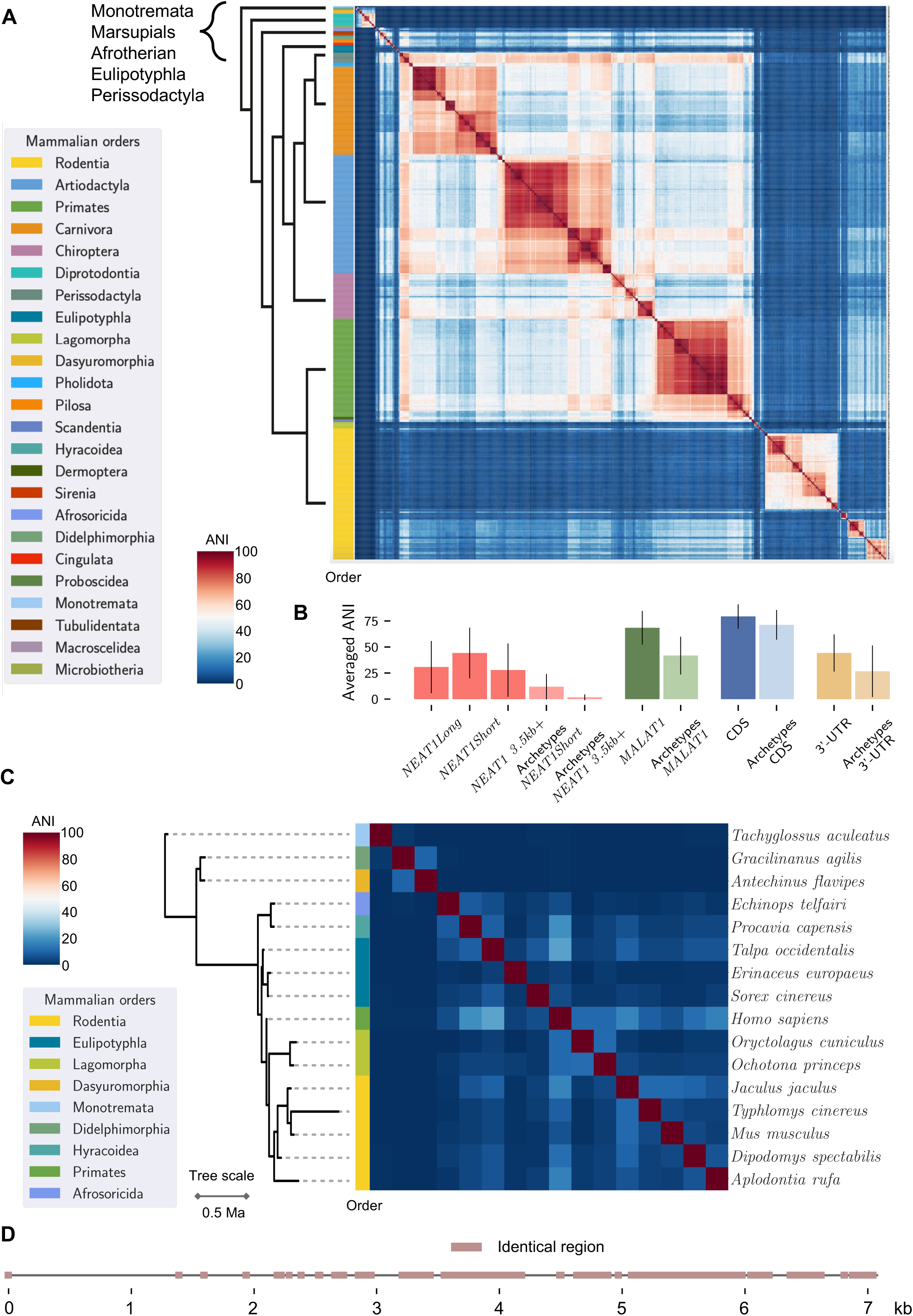
Primary sequence diversity of *NEAT1* and *MALAT1* orthologs in mammals. **A.** Heatmap of ortholog similarity in pairwise comparisons (all-to-all). Orthologs are arranged along a phylogenetic tree, and the colour bar on the left side corresponds to the mammalian orders of individual orthologs; colours are explained in the legend. For visual clarity, the phylogenetic tree was simplified, and information about phylogenetically estimated time was omitted. Red clusters represent groups of highly similar orthologs, while dark blue areas indicate a lack of similarity between them. **B.** Bar plot of averaged ANI for specified groups of orthologs or genes, estimated in pairwise all-to-all comparisons. In addition to the two *NEAT1* isoforms, we also present *NEAT1*_3.5kb+—a part of *NEAT1Long* excluding the 5’-end of the gene, which is shared with *NEAT1Short*. Archetypes refers to a subset of 16 of the most diverged *NEAT1* orthologs. For this species subset, we estimated the average ANI of *MALAT1* orthologs and of protein-coding genes. The averaged ANI of two structural parts of transcripts of protein-coding genes were included for comparison—CDS regions (15,461 orthologous genes were used) and 3’-UTR regions (n=13,847). **C.** Heatmap of primary sequence similarity among *NEAT1* archetypes. Orthologs are arranged along a phylogenetic tree, and the colour bar on the left side corresponds to the mammalian orders of individual orthologs; colours are explained in the legend. The phylogenetic tree is time-scaled. **D.** Regions of identical primary sequence in *MALAT1* orthologs from the archetypes subset (the most diverged sequences, also see Suppl. Fig. 2). Regions are mapped onto human *MALAT1*. ANI = averaged nucleotide identity.

We followed a simple idea: if shared features could be identified in the most divergent sequences of *NEAT1* orthologs, these features would likely have the highest functional relevance. They would also be characteristic of other orthologs, as each divergent sequence would exhibit high homology with some other orthologs, thereby covering the entire diversity of *NEAT1* sequences. Consequently, we identified 16 *NEAT1* orthologs that exhibited the lowest sequence similarity and termed them archetypes (Fig. 4B, C). Some archetypes represented a group of orthologs. For example, human *NEAT1* is a member of the aforementioned largest cluster of orthologs, which we selected as an archetype. Orthologs from the Muridae and Cricetidae families formed the second-largest cluster of *NEAT1* orthologs, and mouse *Neat1* was chosen as an archetype. The remaining archetypes originated from Monotremata, Rodentia (4 archetypes), the Lagomorpha order (2 archetypes), Marsupials (2 archetypes), Eulipotyphla (3 archetypes), Hyracoidea, and the Tenrecidae family (Afrosoricida order) (Fig. 4C). Thus, while some *NEAT1* orthologs exhibited high homology, others showed only traces of sequence similarity and could be used for efficient search of shared features.

*MALAT1* is a more conserved gene than *NEAT1*, with orthologs of Eutherians sharing 60% ANI or higher (Suppl. Fig. 2A, Fig. 4B) and only the orthologs of marsupials and Monotremata were more distinct. Overall, the clustering patterns of heatmaps for both genes were very similar, and the *MALAT1* orthologs in species coding for *NEAT1* archetypes were among the most diverse (Suppl. Fig. 2B, C). We analysed this subset of *MALAT1* orthologs and identified positions in multiple sequence alignments that were identical among the archetypes. These identical regions covered around 13% of the *MALAT1* sequence (Fig. 4D) and were mostly distributed in the 3’-end of the gene. These findings suggest a high functional importance of the primary sequence of *MALAT1*, particularly its 3’-end.

To estimate the degree of *NEAT1* and *MALAT1* sequence variation, we compared the averaged ANI of these genes to the averaged ANI of CDSs and 3’-UTRs of transcripts of orthologs of protein-coding genes in mammals (Fig. 4B). In this analysis, we included the *NEAT1Short* isoform and the part of *NEAT1Long* after the polyadenylation signal (*NEAT1*_*3.5kb+*). The results showed that *MALAT1* was close to CDSs in terms of conservation, while *NEAT1* was similar to 3’-UTRs when considering all full-length orthologs. Notably, *NEAT1Short* is significantly more conserved than the *NEAT1_3.5kb+* region, and the *NEAT1Short* isoform exhibited some sequence similarity among archetypes, whereas similarity in the *NEAT1_3.5kb+* region was nearly absent. This higher level of conservation, comparable to that of 3’-UTRs, underscores the potential functional importance of the primary or secondary structure of the *NEAT1Short* isoform. A similar conclusion can be drawn for *MALAT1* due to its exceptional level of conservation, comparable to CDSs, while the primary sequence of *NEAT1_3.5kb+* appears to be highly variable.

### Transposable elements contribute to the diversity of *NEAT1*

Due to the high diversity of the primary sequences of *NEAT1* orthologs, we focused on identifying shared features that could be detected without the use of multiple sequence alignment. First, we examined the contribution of transposable elements (TEs) to the diversity of *NEAT1* orthologs. We predicted non-overlapping positions of TEs in all orthologs of *NEAT1* and *MALAT1*. In human *NEAT1*, we found four large fragments of LINE elements and six complete SINEs. These six SINEs included four Alu elements and two FLAM-C elements. Two of the identified Alu elements, AluSx3 at 17,804-18,067bp and AluJr at 17,532-17,678bp, can form an IRAlu secondary stem loop structure, which may attract ADARs for A-to-I modification of *NEAT1* (Vlachogiannis et al., 2021). In mouse *NEAT1*, we identified four SINE elements (types B1_Mus1, B3, B1_Mm, and B1_Mus2), which are non-complementary to each other.

Analysis of TEs in *NEAT1* orthologs revealed their high diversity and enrichment (Fig. 5A). Among the predicted TE types, SINEs and LINEs were predominant, with varying frequencies across different taxa. Although ERVs are known as the most common type of TEs in lncRNAs (Kelley and Rinn, 2012), we identified only 15 cases of ERV integration in *NEAT1* orthologs from seven different orders (Suppl. Fig. 3A). We also did not find any distinguishable pattern in the distribution of TEs across archetypes (Suppl. Fig. 3B). While our TE prediction method depends on how well TEs are studied in specific groups of mammals—which may affect the identification of exact TE types and frequencies—we can still gain a general impression of the importance of TEs in the evolution of *NEAT1*.

**Figure 5.**
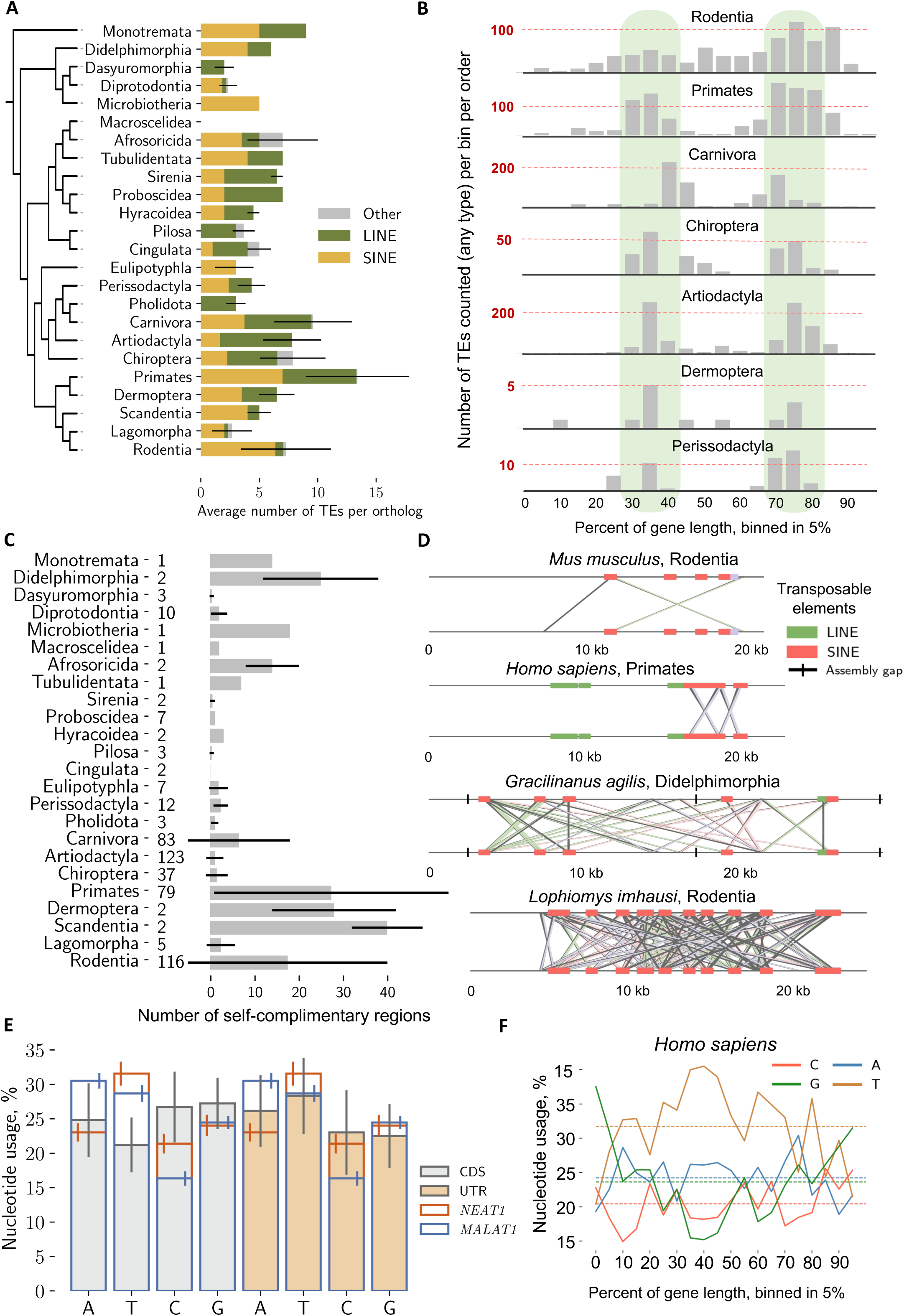
Transposable elements contribute to *NEAT1* sequence diversity. **A.** Bar plot of the average number of TEs per ortholog across mammalian orders. The phylogenetic tree is not time-scaled. **B.** Distribution of TEs within *NEAT1* orthologs. The length of individual orthologs was binned into 5% segments, and the number of annotated TEs within each bin was summed for all orthologs per mammalian order. Two segments (30-40% and 70-80% bins), which most frequently contain TEs, are highlighted in green. **C.** Bar plot showing the number of self-complementary regions per ortholog, averaged per mammalian order. The number of orthologs used in the assessment is indicated on the left. **D.** Graphical representation of the distribution of self-complementary regions in four selected *NEAT1* orthologs. Each ortholog is aligned to itself, and reverse complementary regions are connected with lines, forming a visual ‘cross’ shape. The second ‘cross’ (around 20kb) in human *NEAT1* corresponds to an IRAlu-formed hairpin. Self-complementary regions often coincide with TEs and can occur over short distances, resembling the IRAlu of human *NEAT1*, or over much longer distances. Additional plots are in Suppl. Fig. 3C. **E.** Averaged nucleotide usage (nucleotide composition of a molecule estimated as a percentage) of *NEAT1* and *MALAT1* orthologs compared to the averaged nucleotide usage of CDS and 3’-UTRs of protein-coding genes. The nucleotide usage of *NEAT1* and *MALAT1* is overlaid on coloured bars representing the nucleotide usage of CDSs (grey bars) and 3’-UTRs (orange bars). The standard deviation whisker is shifted for visual clarity. **F.** Plot of nucleotide usage of human *NEAT1* along the sequence. The length of the ortholog was binned into 5% segments, and the nucleotide usage of each bin was estimated. The average nucleotide usage of human *NEAT1* is depicted with dashed lines. Additional plots for *NEAT1* are in Suppl. Fig. 6, and for *MALAT1* in Suppl. Fig. 7.

As the next step, we analysed the integration positions of TEs in *NEAT1* orthologs. We divided the length of each ortholog into 5% bins, counted the number of TEs in each bin, and summed them per taxon (Fig. 5B). With this approach, we found that a few taxa exhibited a bimodal distribution of integration sites, around 30-40% and 70-80% of the gene length. These orders included Carnivora, Artiodactyla, Primates, and Chiroptera. However, in Rodentia, TEs were broadly distributed, with a slight preference for the end of the gene (Fig. 5B). Considering the high diversity of TEs in *NEAT1* orthologs, our results suggest that these two regions can tolerate TE integration without affecting function.

Regions of self-complementarity can potentially contribute to secondary structure formation and paraspeckle stabilisation. For example, IRAlu elements, which are regions of self-complementarity in close proximity, can form stem loops that contribute to *NEAT1* A-to-I modification and paraspeckle assembly via interaction with NONO and SFPQ (Vlachogiannis et al., 2021), (Knott et al., 2016). We studied the presence of self-complementary regions in *NEAT1* and *MALAT1* orthologs and found that these regions were common in *NEAT1* but not in *MALAT1* (Fig. 5C, D, Suppl. Fig. 3C). Specifically, we identified self-complementary regions in 71% of *NEAT1* orthologs, with 14.68 ± 20.85 regions per ortholog, and *Lophiomys imhausi* (Rodentia) exhibiting the maximum recorded number of 132 regions (Fig. 5D). We observed that some of these interactions occurred over long distances, while others were in close proximity, potentially mimicking the IRAlu situation (Fig. 5D, Suppl. Fig. 3C). This diversity of interactions could be explained by the bimodal pattern of TE distribution, as we also noted that TEs were frequently the sources of these complementary regions.

TEs were rarely localised within *NEAT1Short* isoforms. We identified only 49 cases in six mammalian orders: most of them from Rodentia (18 orthologs), the Canidae family of Carnivora (15), Primates (11, with six in the Lemuridae family), Sirenia (2), Afrosoricida (2), and one from Dermoptera (Suppl. Fig. 4A).

It has been shown that mouse *Malat1* contains the SINE B2 element. However, this is an exception; we noted only 13 orthologs in our collection with a single TE. The Rodentia order included 12 of these: all species in the *Mus* genus contained B2 SINEs, some species of the Sciuridae family had *de novo* identified DNA elements, and the naked mole-rat possessed a B4A SINE. Another case was found in *Ailurus fulgens* (Carnivora). All these TEs were localised in close proximity to the 5ʹ-end (Suppl. Fig. 4B). Therefore, *MALAT1* is rarely affected by TE activity.

As SINEs typically integrate into A-T enriched areas (Daniels and Deininger, 1985), we analysed nucleotide usage in *NEAT1* and *MALAT1* orthologs to gain mechanistic insight (Suppl. Fig. 5). We found a high enrichment of T and a depletion of C nucleotides in almost all orthologs of both genes. *MALAT1* orthologs additionally exhibited a high proportion of A nucleotides, demonstrating a nucleotide composition more prone to TE integration (Fig. 5E, Suppl. Fig. 5). To determine whether these patterns are common to other genes, we compared them to CDS and 3ʹ-UTR regions of protein-coding genes in mammals (Fig. 5E). This analysis showed enrichment of C and G nucleotides in CDSs and A and T nucleotides in 3ʹ-UTRs. Additionally, it has been shown that 3ʹ-UTRs are also prone to TE integration (Lagemaat et al., 2003), which aligns well with the nucleotide usage profile. *NEAT1* and *MALAT1* were similar in composition to 3ʹ-UTRs (genes were within the standard deviation), although *MALAT1* exhibited an even stronger depletion of C nucleotides. This indicates that from a sequence composition perspective, *MALAT1* exhibited an exceptionally low TE frequency.

Finally, we analysed nucleotide usage along the sequences of the two genes. We identified peaks of G nucleotide usage at both ends of the *NEAT1* gene, with a more pronounced peak at the 5ʹ-end (Fig. 5F). This pattern was noticeable in almost all archetypes (Suppl. Fig. 6). Overall, the A-T enriched central part of *NEAT1* coincided well with the hot spots of TE integration. In *MALAT1* orthologs, the nucleotide usage pattern differed, showing a peak of A nucleotide usage at the 5ʹ-end of the gene, which correlates with the integration sites of the infrequently detected TEs (Suppl. Fig. 7). To summarise, we found high frequency of TE’s integration in specific regions of *NEAT1Long*, which corresponds well to sequence composition and introduces self-complementary interactions. In contrast, TE integration was exceptionally low in *NEAT1Short* isoforms and in *MALAT1* orthologs.

### G-quadruplexes and binding sites for TDP-43 are common features in archetypes

The next group of features we focused on in our analysis was the examination of short primary sequence patterns. Guanine tracks separated by loops can form G-quadruplexes— secondary structures which, in human *NEAT1* and *MALAT1*, facilitate interactions with NONO ((Arun et al., 2020), (Mou et al., 2022)). We predicted G-quadruplexes in *NEAT1* orthologs and identified them in all orthologs, including all archetypes, with an average of 19.2 ± 5.9 per ortholog. In the archetypes, they predominantly localised at both ends, within the ‘shell’ area of the paraspeckles (Fig. 6A). In archetypes they predominantly localised at both ends, within the ‘shell’ area of the paraspeckles (Fig. 6A). This observation aligns well with our finding of shifts in nucleotide usage at both ends of *NEAT1*, showing enrichment in G nucleotides. G-quadruplexes were also typical for *MALAT1* orthologs, though in much lower numbers (9.1 ± 1.6 per ortholog). We compared our results to the frequencies of G-quadruplexes in CDSs and 3’-UTRs of orthologs of protein-coding genes in mammals (Fig. 6B) and found that the number of G-quadruplexes in *NEAT1* orthologs was exceptionally high. Additionally, the abundance of G-quadruplexes in *MALAT1* orthologs was also very high and more characteristic of CDSs rather than 3’-UTRs. Our findings point to the significant functional importance of G-quadruplexes in both genes.

**Figure 6.**
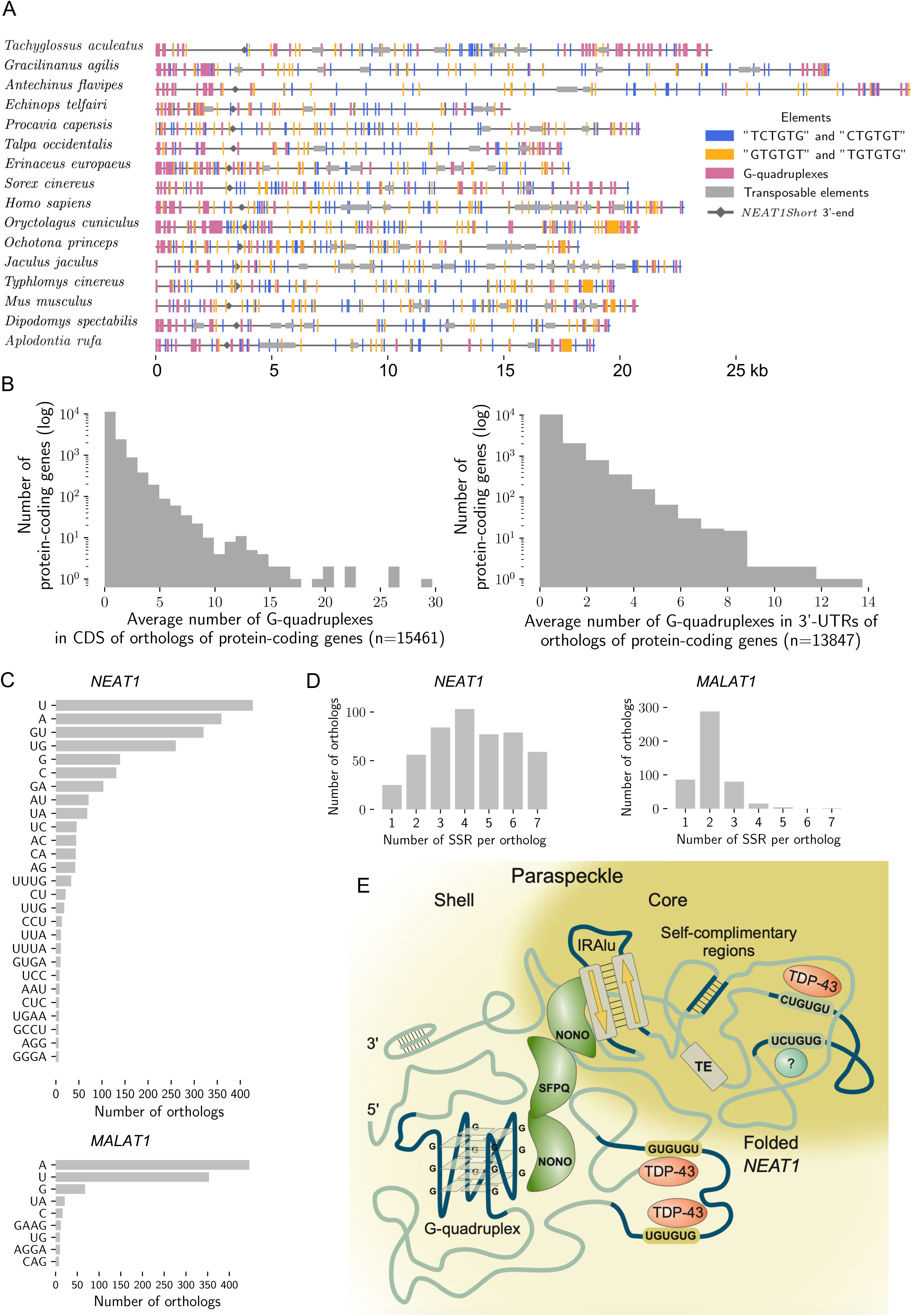
Simple and complex repeats in *NEAT1* and *MALAT1* orthologs. **A.** Distribution of two groups of hexamers and G-quadruplexes in *NEAT1* archetypes. Each vertical line, in the colours described in the legend, marks the position of the identified element. **B.** Frequency of G-quadruplex detection in CDSs and 3’-UTRs of protein-coding transcripts. The number of detected G-quadruplexes in orthologs was averaged per gene and used in the plot. **C.** Frequency of SSR detection in *NEAT1* and *MALAT1* orthologs. Only the most frequent SSRs are shown for *NEAT1*. To be counted as SSRs, an individual nucleotide should be repeated at least 10 times. **D.** Number of SSRs per *NEAT1* and *MALAT1* ortholog. **E.** Summary of identified conserved features of *NEAT1*, contributing to the function and stabilisation of paraspeckles. Most of the features show a preference for predominant localisation in the ‘shell’ or ‘core’ parts of the paraspeckle. G-quadruplexes and IRAlu (and possibly closely located self-complementary regions) serve as binding sites for NONO and likely other proteins of the DSHS family, along with many other resident proteins. NONO, together with other multidomain proteins, cross-stitches *NEAT1*, stabilising the paraspeckle. Long-range self-complementary regions can play the same role. GU-repeats may be bound by TDP-43, which is predominantly located in the ‘shell’ part of the paraspeckle.

The subset of *NEAT1* archetypes presents an opportunity to identify frequent or systematically recurring sequence motifs that are universally important for paraspeckle formation and function. We chose hexamers as an optimum between diversity and uniqueness, given that the 4,096 possible combinations of letters in hexamers are theoretically diverse enough to appear only once or twice in the longest *NEAT1* ortholog, which contains 6,075 hexamers. Longer motifs are more diverse and unique, making it less likely to find them across all orthologs.

As a result of hexamer profiling, we identified two groups of motifs that are both frequent and shared by all *NEAT1* archetypes. The first group comprised ‘GU’-based hexamers (’GUGUGU’ and ‘UGUGUG’), which are known TDP-43 binding sites (Rot et al., 2017). These hexamers were primarily localised at the gene ends, in the ‘shell’ region of paraspeckles, which is also enriched in TDP-43 (West et al., 2016) (Fig. 6A). The second group of motifs included ‘UCUGUG’ and ‘CUGUGU’, which displayed a different distribution pattern, frequently localising in the central part of *NEAT1*, the ‘core’ of paraspeckles. These motifs may also be recognised by TDP-43 (Rot et al., 2017); however, the difference in distribution patterns suggests distinct regulatory mechanisms and possibly varying binding affinities for TDP-43. Additionally, these motifs may serve as binding sites for alternative proteins. Although the distribution of identified hexamers and G-quadruplexes did not show a strong association with TE integration sites (Fig. 6A), some of these elements were located within predicted TEs.

We observed that different archetypes had varying most abundant hexamers (Suppl. Table 1). In some archetypes, these hexamers clustered and formed areas of low sequence complexity (Suppl. Fig. 8). However, the distribution of these clusters was archetype-specific and did not exhibit any common pattern, therefore not suggesting any specific functional role.

Finally, we analysed simple sequence repeats (SSRs, Fig. 6C, D). This type of analysis requires that a single nucleotide is repeated at least 10 times and a two-nucleotide pattern at least 6 times, which differs from hexamer profiling. We noted that *NEAT1* orthologs contained on average 3-4 SSRs, while in *MALAT1*, SSRs were less abundant (Fig. 6D). The diversity of SSRs in *NEAT1* was also higher. While in *MALAT1*, SSRs comprised stretches of A or T nucleotides, in *NEAT1*, ‘GU’ (and ‘UG’) repeats were among the most abundant and could be found in approximately half of the orthologs (Fig. 6C). This observation aligns well with the hexamer profiling results.

Overall, using the subset of *NEAT1* archetypes, we identified G-quadruplexes, TDP-43 binding motifs, and a diversity of self-complementary regions as shared features important for paraspeckle function. This highlights the mechanism by which, in the absence of primary sequence similarity, function can be maintained through securing interactions with proteins essential for paraspeckle assembly. A summary of all the critical findings is presented in Figure 6E.

### Taxa-specific speed of *NEAT1* evolution

By analysing the subset of archetypes and scrutinising the heatmap of all-to-all *NEAT1* ortholog similarity (Fig. 4A), we noted that sequence diversity among orthologs varied between taxa. For example, orthologs of Carnivora and Artiodactyla formed distinct clusters (Fig. 4A), where they were highly similar to each other within the taxon (60-70% ANI) and also between the taxa (40-50% ANI). However, several archetypes—sequences with ANI lower than 10% to any other ortholog—originated from the orders Rodentia, Eulipotyphla, or Lagomorpha, demonstrating that even within a taxon, orthologs can be highly divergent. Differences between orthologs from different mammalian orders cannot always be explained by the phylogenetic tree and the evolutionary time since the taxa diverged. For example, Rodentia and Lagomorpha are phylogenetically closer to Primates than to Carnivora, yet orthologs of Primates are much more similar to those of Carnivora than to Rodentia and Lagomorpha (Fig. 4A). Therefore, we decided to explore the evolutionary dynamics of *NEAT1* further.

We focused on the Rodentia order, as it comprised the highest number of sequenced genomes and identified orthologs, and exhibited a high diversity in their primary sequences (Fig. 7A, B). Some of the clusters distinguished in the heatmap aligned well with the borders of individual families, with reduced similarity between them (Fig. 7A). However, other clusters did not. One of these was a cluster showing high similarity among orthologs from the Muridae and Cricetidae families. Another was a cluster with high similarity among orthologs from 13 different families (red dashed frame in Fig. 7A). Thus, overall taxonomic borders within the Rodentia order did not provide a strong explanation for *NEAT1* sequence variability.

**Figure 7.**
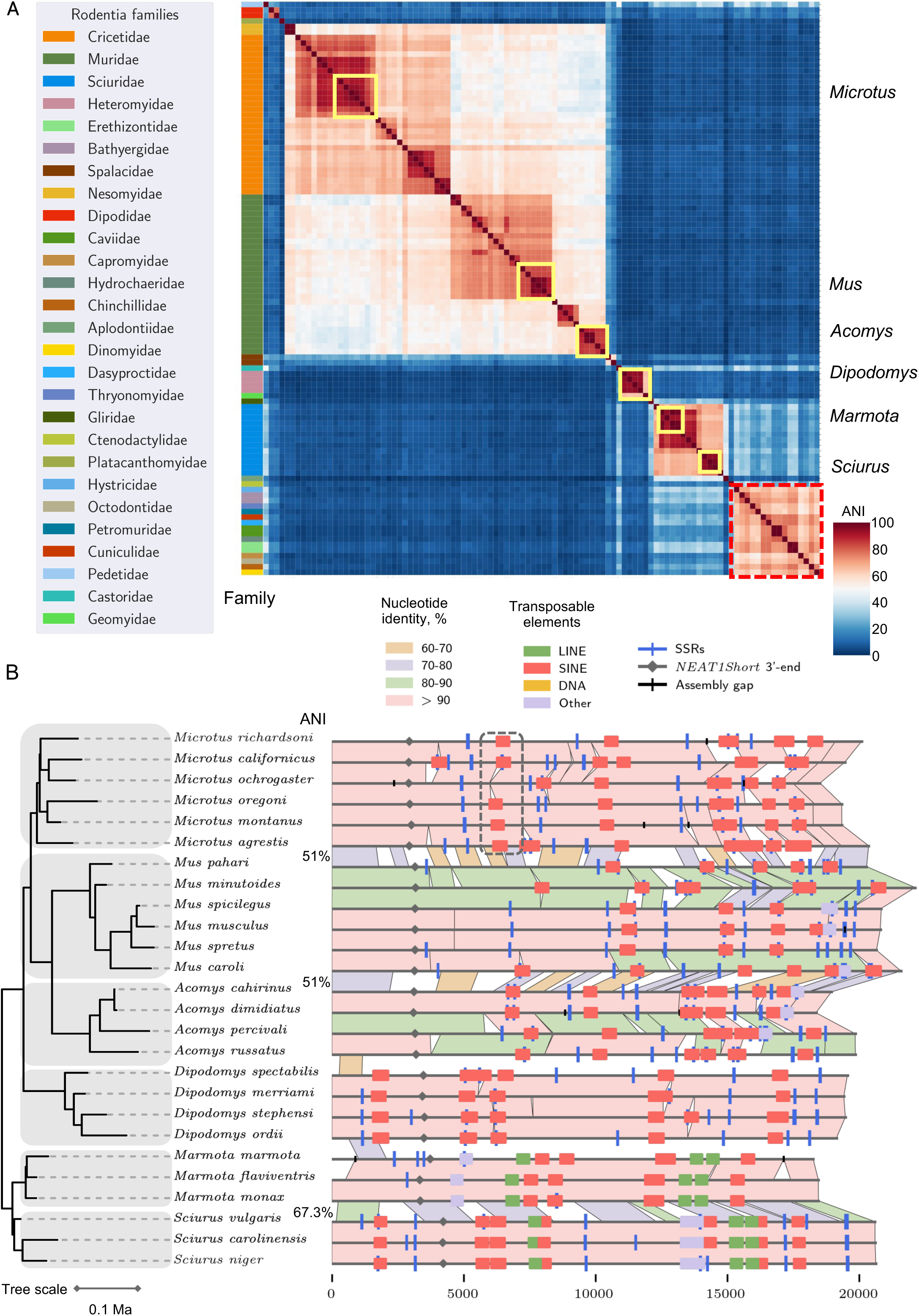
Uneven speed of *NEAT1* evolution. **A.** Heatmap of primary sequence similarity between *NEAT1* orthologs of the Rodentia order, in pairwise all-to-all comparison. Clusters of red represent groups of highly similar orthologs, while dark blue areas indicate a lack of similarity between them. The colour bar on the left represents individual families, as seen in the legend. Six selected genera for the alignment in part B are highlighted with yellow frames. The red dashed frame highlights the similarity cluster of 13 Rodentia families mentioned in the Results section. **B.** Graphical representation of the pairwise alignment of *NEAT1* orthologs from six Rodentia genera. Orthologs are aligned according to the time-scaled phylogenetic tree on the left. Individual genera are highlighted in grey for visual clarity. *Mus* and *Acomys* genera exhibit a higher evolutionary rate compared to the others. An example of an excised SINE element is highlighted with a dashed frame.

As a next step, we analysed the evolutionary dynamics at the genus level. We selected six Rodentia genera with orthologs from at least three species and performed pairwise alignments along the phylogenetic tree (Fig. 7B). The highest primary sequence variation was detected in the genera *Mus* and *Acomys*, while other genera exhibited relatively high levels of conservation. This difference in the rate of evolution was also visible between the genera. Specifically, the evolutionary divergence of the genera *Sciurus* and *Marmota* occurred earlier than that of *Mus*, *Acomys*, and *Microtus*, yet the similarity of orthologs between these taxa is the highest (67.3% ANI, Fig. 7B), highlighting their high degree of conservation. The divergence between *Mus* and *Acomys* occurred later than between either of them and *Microtus*, although the similarity level was the same (51% ANI) between *Mus* and *Acomys* and between *Acomys* and *Microtus*, suggesting a high rate of evolution in *Mus* and *Acomys* and a higher conservation level in *Microtus*. Altogether, these patterns indicate that different taxa may have been subjected to varying evolutionary pressures.

The activity of TEs in *NEAT1* orthologs was clearly one of the main contributors to sequence variation, where their integration or deletion caused local sequence changes. The developed graphical representation of pairwise alignments made this dynamic visible (Fig. 7B, Suppl. Fig. 9, 10). Special cases illustrating the importance of TEs in shaping *NEAT1* sequence variability included those originating from several assemblies of the same species (Suppl. Fig. 9). In these cases, sequence divergence was minimal, and only TE integration events accounted for the differences. One key difference between SINE and LINE elements is that SINEs depend on LINEs for amplification. Moreover, a specific mechanism for SINE excision has not been identified (Batzer and Deininger, 2002), suggesting that SINEs remain at their integration site and disappear as part of a mutational process (Richardson et al., 2015). However, we identified three instances of SINE element excision. For example, in Figure 7A, in the highlighted area (and Suppl. Fig. 10), a SINE shared by the entire *Microtus* genus was excised in *Microtus ochrogaster*. This observation suggests the existence of a mechanism for SINE excision and that, overall, TE dynamics is one of the major factors shaping *NEAT1* evolution.

## Discussion

Here, we used a phylogenetic approach to identify the basis for *NEAT1* and *MALAT1* function. A particular advantage of our research was the high degree of conservation in the 3’-end structures of *NEAT1* and its promoter region, which made it possible to identify *NEAT1* orthologs across diverse mammals, regardless of the homology level in other gene regions. It is widely recognised that signs of purifying selection serve as hallmarks of functional importance, manifesting in the conservation of sequence and/or structural elements across a significant evolutionary distance. The selection of the archetypes subset created a unique opportunity to decipher the structure-function relationship of *NEAT1*—one of the diverse lncRNAs, which exhibited an overall conservation level lower than that of UTRs. The *NEAT1* gene gives rise to at least three sub-molecules—*NEAT1Long*, *NEAT1Short*, and a tRNA-like structure—each characterised by different conservation levels, pointing to distinct functional roles, which we discuss separately.

### Architectural long NEAT1 isoform

The long isoform of *NEAT1* forms the backbone of paraspeckles, seeding and stabilising their assembly through interactions with multidomain proteins. We summarised our findings of the conserved structural elements of *NEAT1Long* in Figure 6E. Most of these elements are involved in maintaining paraspeckle integrity, providing interaction sites for multidomain proteins, and facilitating interactions with TDP-43—a protein crucial for regulating *NEAT1* isoform switching.

The length of RNA in condensate formation is a critical parameter: longer molecules enable faster and easier aggregation, as demonstrated in experimental conditions using synthetic RNAs (reviewed in (Garcia-Jove Navarro et al., 2019), (Van Treeck and Parker, 2018)). Furthermore, testing the length of RNA molecules isolated from stress granules in the human U-2 cell line revealed that they were significantly longer than RNAs dissolved in the cytoplasm (Khong et al., 2017). This aligns with the exceptional length of *NEAT1* orthologs, the shortest of which totals 14.5 kb. This remarkable length, which facilitates self-aggregation, may also explain why paraspeckle formation begins immediately at the transcription site, with multidomain stabilising proteins recruited later (Mao et al., 2011). The substantial variation in the length of *NEAT1* orthologs raises an open question about the potential variation in the physical properties of paraspeckles across species, such as differences in the speed of paraspeckle assembly, their linear size, stiffness, and penetrability.

We have previously shown that the isoform switch of human and murine *NEAT1* is regulated by TDP-43 through binding to GU repeats, which abrogate the polyadenylation of the short isoform (Modic et al., 2019). In the same study, we also demonstrated that GU repeats are the predominant mechanism for TDP-43 sequestration in paraspeckles. Here, we noted that the presence of GU repeats is a universal feature of *NEAT1* archetypes, predominantly localised in areas closer to the 5’ and 3’ ends. This observation aligns well with the fact that TDP-43 is more frequently detected in the shell area of paraspeckles (West et al., 2016). Thus, the interaction between *NEAT1* and TDP-43 is clearly one of the crucial functional aspects of paraspeckles, and the mechanism of isoform switching via TDP-43 association with GU repeats in *NEAT1* is likely conserved across all mammals.

G-quadruplexes are another universal feature of all *NEAT1* orthologs. The importance of these structures in the development of various diseases, including cancer and neurodegenerative pathologies, has become evident over the past decade (Cammas and Millevoi, 2017; Wang et al., 2021). These structures can spontaneously form in both RNA and DNA. Functionally, G-quadruplexes are involved in almost all aspects of gene expression regulation, from transcription to translation, in the modification of mRNAs and miRNAs, and are also associated with phase separation processes (Asamitsu and Shioda, 2021; Dumas et al., 2021). Moreover, RNA stretches of guanine nucleotides, which can spontaneously form G-quadruplexes, can facilitate the formation of condensates (Van Treeck and Parker, 2018). It has been shown that NONO can interact with the G-quadruplexes of both *NEAT1* and *MALAT1* genes (Arun et al., 2020), (Mou et al., 2022). However, if we compare the list of RNA-binding proteins capable of interacting with G-quadruplexes (Bourdon et al., 2023) to the list of resident proteins in paraspeckles (Fox et al., 2018), we observe a significant overlap. For example, among the essential paraspeckle proteins—HNRNPH3, HNRNPK, RBM14, SMARCA4, NONO, SFPQ, and TDP-43—along with 17 other non-essential proteins, including another DBHS family protein -PSPC1, are all capable of binding to G-quadruplexes. The diversity of paraspeckle proteins that recognise G-quadruplexes offers the potential for interchange between them in maintaining paraspeckle integrity, which may explain the importance but non-essentiality of certain proteins (Fox et al., 2018). This possibility of exchange was demonstrated by Yamada *et al* (Yamada et al., 2022) for the FET family of proteins. They showed that, in tissues of the naked mole-rat containing intact paraspeckles, the distal part of the intestinal epithelium lacked the essential protein FUS, while expressing other proteins of the same family, EWSR1 and TAF15. Overall, G-quadruplexes provide a potential mechanism for the recruitment of mRNAs, miRNAs, and proteins to paraspeckles, as well as facilitating interactions with DNA, where G-quadruplex-binding proteins can cross-stitch paraspeckles to DNA.

Self-complementary interactions between inverted repeat Alu elements (IRAlu) in humans can form stem loop secondary structure, attracting ADAR enzymes to perform A-to-I RNA modification. Specifically, IRAlu structures have been identified in human *NEAT1*, triggering the modification of adenines in their proximity (Vlachogiannis et al., 2021). Additionally, IRAlu elements can be recognised and bound by NONO (Elbarbary and Maquat, 2015). Although Alu elements are typical only for primates, we found that many *NEAT1* orthologs are characterised by reverse complementary regions, frequently originating from diverse TEs. We speculate that these regions, being in close proximity to each other, may also have the potential to form stem loop structures, which could potentially be recognised by NONO and/or ADAR enzymes. Moreover, when located farther apart, these self-complementary regions could contribute to paraspeckle stabilisation, particularly in the early stages of paraspeckle assembly before the recruitment of multidomain proteins.

### Short isoform of NEAT1

Our research highlights the distinctive properties of *NEAT1Short*. First, we predicted its presence in all mammalian orthologs, emphasising its universality. *NEAT1Short* is subject to much stronger purifying selection than *NEAT1Long*. The conservation of *NEAT1Short* isoform length is also very high, and it does not have a proportional relationship to the length of *NEAT1Long*. We detected only a small number of cases where *NEAT1Short* contained TEs, and overall, *NEAT1Short* was depleted of both simple and more complex repeats. These findings indicate a distinct functional trajectory for *NEAT1Short*, separate from *NEAT1Long*, about which there is currently limited knowledge. For example, it has been shown that *NEAT1Short* can be located outside of paraspeckles and concentrated in much smaller foci known as ‘microspeckles,’ the function of which remains unclear (Li et al., 2017). In experiments conducted by Naveed et al., it was demonstrated that *NEAT1Short* can have an effect on cell proliferation that is opposite to that of *NEAT1Long* (Naveed et al., 2021). Altogether, we believe it is urgently necessary to begin unravelling the function of *NEAT1Short* and to delineate the functional differences between the two isoforms in experimental designs.

### tRNA-like structure

The primary sequence of the tRNA-like structure of *NEAT1* varies more than that of *MALAT1* (mascRNA), although, overall, they remain highly similar. This may indicate functional differences between these two structures. First, the tRNA-like structure is crucial for the maturation processes of both genes and must be effectively recognised by RNase P and Z. Therefore, the most conserved parts shared by both structures likely serve this role, which includes the tRNA-like conformation itself and the highly conserved hairpin III (Fig. 2A,B).

It has been shown that mascRNA may additionally play a role in cellular metabolism within the cytoplasm. For example, it can contribute to increased protein translation and cell proliferation by binding to the multi-tRNA synthetase complex (Lu et al., 2020), or it may influence immune response by impacting macrophages (Gast et al., 2022). Dissimilarly, NEAT1’s tRNA-like molecules were shown to degraded in cell lines. Based on these differences it is important to systematically analyse the functions of tRNA-like molecules in different cell types and animals as they may have been adapted to perform specific functions.

### From conserved transcriptional regulation to NEAT1’s role in cell biogenesis

We broadened our investigation of the structure-function axis of *NEAT1* with a functional analysis of conserved TFs motifs identified in the promoter regions of *NEAT1* orthologs. This provided an overview of the biological processes in which the gene is involved, many of which already have extensive experimental confirmation. *NEAT1*’s involvement in apoptosis and proliferation is well established. It is a downstream gene of p53 (Adriaens et al., 2016) and is also involved in regulating the availability of the 53BP1 protein, which activates p53 in response to DNA damage (Kilgas et al., 2024). *NEAT1* expression and the number of paraspeckles are also upregulated in response to various stresses (McCluggage and Fox, 2021). The association of *NEAT1* with diverse neurodegenerative diseases is well documented (An et al., 2018), and the identification of TFs involved in CNS development further expands *NEAT1*’s role in nervous system biogenesis.

The number of paraspeckles (and *NEAT1* expression levels) oscillates with circadian rhythms, releasing IRAlu-containing mRNAs (Torres et al., 2017, 2016) and regulating 53BP1 availability in a cell-cycle-dependent manner (Kilgas et al., 2024). Moreover, *NEAT1* directly binds approximately 30% of all mRNAs located in paraspeckles, most of which are also involved in circadian rhythm cycles (Jacq et al., 2021). Additionally, NONO and SFPQ are known to be involved in circadian rhythm regulation (Knott et al., 2016; Kowalska et al., 2012). Our analysis also suggests a potential role for *NEAT1* and *MALAT1* in spermatogenesis and gonad development, although this has not yet been experimentally confirmed.

### NEAT1 and MALAT1: uniquely similar but different lncRNAs

Our study confirms the synteny of *NEAT1* and *MALAT1* across the full range of mammalian species: these genes are located in close proximity on the same strand. The uniqueness and similarity of their gene maturation processes, both utilising tRNA processing machinery, along with their roles in spatially associated nuclear bodies, raise the expectation of similar regulation, conservation, and function for *NEAT1* and *MALAT1*. However, this is not the case: *MALAT1* is a highly conserved lncRNA, while *NEAT1* is more variable.

This difference in conservation is possibly associated with the frequency of TEs integration, as *NEAT1* is more prone to such integrations compared to *MALAT1*. However, our analysis of nucleotide usage highlighted an opposite trend: *MALAT1* has, on average, a more favourable nucleotide composition for TE integration. This further underscores the functional importance of conserved primary sequence of *MALAT1*. Our results indicated that the central part of *MALAT1* and its 3’-end exhibit the highest levels of conservation, whereas the more variable 5’-end can carry inserted TEs. Although there is a large body of research on the role of *MALAT1* in various pathological conditions (such as cancer), the exact mechanisms of its action remain unclear. It has been shown that two regions in *MALAT1*, located approximately at 2-3 kb and 6-7 kb, are responsible for its localisation in nuclear speckles (Miyagawa et al., 2012), which aligns with our results showing a high level of sequence conservation in these regions. Accumulation of mutations in another conserved region of *MALAT1* (3-4.3 kb) has been associated with breast cancer progression (Ellis et al., 2012). Together, these findings suggest that *MALAT1*’s primary sequence plays a major role in its function, while for *NEAT1*, secondary structural elements appear to be more crucial.

The analysis of the conservation of promoter regions, TATA-boxes, and transcription TF binding sites revealed another key difference between *NEAT1* and *MALAT1*. Although *MALAT1* showed greater gene conservation than *NEAT1*, the variability in *MALAT1*’s promoter region and potential transcriptional regulation was higher. This suggests that *MALAT1* may have adapted to different gene networks across species, while *NEAT1* remains a consistent player in the same biological processes.

### Uneven speed of NEAT1 evolution

Our research identified two main mechanisms driving *NEAT1* evolution: divergence due to the accumulation of mutations and the high frequency of TEs integration and excision. TEs not only lead to the loss of primary sequence regions but also influence gene length in both directions—making it longer through integration or shorter through excision—explaining the considerable variation in gene length across mammals. TEs also introduce self-complementary regions, stabilising paraspeckles, as well as repeats and G-quadruplexes, which serve as interaction sites for key resident proteins. Therefore, TEs play a crucial role in *NEAT1* adaptation overall.

Our analysis highlighted taxa with accelerated *NEAT1* evolution, such as Eulipotyphla, Lagomorpha, and the *Mus* and *Acomys* genera of the Rodentia order. This phenomenon of varied evolutionary speed has been previously demonstrated for some lncRNAs. For example, unannotated and largely non-coding human accelerated regions (Pollard et al., 2006b, 2006a) are conserved genomic regions across mammals that accumulate disproportionately more mutations in humans, many of which function as enhancers in neurodevelopment (Doan et al., 2016; Girskis et al., 2021). In this case, the rate of evolution highlights species or taxon-specific adaptations to their ecological niches. We speculate that this mechanism may also influence *NEAT1* biogenesis, as *NEAT1* can directly interact with diverse mRNAs and miRNAs, possibly via complementary interactions of primary sequences. This may explain the high evolutionary speed observed in certain taxa and across mammals in general.

Hezroni et al. identified many lncRNAs with conserved synteny across vertebrates, which exhibit sequence homology only among closely related species while lacking similarity over greater phylogenetic distances (Hezroni et al., 2015). Our study provides possible explanations of functions of such lncRNAs. Specifically, interactions with functionally crucial partners may be mediated by short secondary structure elements or short sequence-specific recognition sites, which are not easily detectable by traditional homology search algorithms.

## Conclusions

To gain critical insights into the sequence–structure–function axis of *NEAT1* and *MALAT1*, we utilised a comparative phylogenetic approach. We identified evidence of purifying selection in features such as the tRNA-like structures and triple helices of both lncRNAs, G-quadruplexes and TDP-43 binding motifs in *NEAT1*, alongside the overall high conservation of *MALAT1*’s primary sequence. The universality of the *NEAT1Short* isoform, its higher conservation level in terms of primary sequence and length, suggests a distinct functional role for *NEAT1Short*. Our findings also support the potential conservation of the isoform switch mechanism mediated by TDP-43. We demonstrated the crucial role of TEs in *NEAT1* evolution, influencing gene length, introducing stabilising self-complementary regions and stem loop structures possibly recognised by NONO, and contributing to the delivery of new G-quadruplexes and short protein-binding motifs.

## Methods

### Identification of coordinates for *NEAT1* and *MALAT1* orthologs

Mammalian genomes were downloaded from GenBank ((Clark et al., 2016), July 2023). Annotated *NEAT1* and *MALAT1* orthologs from *Homo sapiens* (NR_131012.1), *Mus musculus* (NR_131212.1,(O’Leary et al., 2016)), and *Monodelphis domestica* (KX036207.1, (Cornelis et al., 2016b)) were used for similarity searches and the identification of orthologs in the downloaded genomes. We additionally retrieved promoter regions and triple helix motifs, followed by tRNA-like structure sequences, for these annotated orthologs using in-house scripts. These sequences were subjected to a blastn (Altschul et al., 1990) search against the downloaded mammalian genomes. Approximate gene coordinates were obtained from the homology search results and were complemented with some manual curation in cases where *NEAT1* and *MALAT1* orthologs were found on different contigs due to fragmentary assembly. Genes were retrieved with some sequence excess at both the 5’-and 3’-ends and subjected to multiple sequence alignment (MSA, MAFFT, v7.487, (Katoh and Standley, 2013), default parameters). Since *NEAT1* showed noticeably higher divergence compared to *MALAT1*, we divided the mammals into eight groups according to the phylogenetic tree ((Ns et al., 2019)).

Group1: Monotremata, Didelphimorphia, Microbiotheria, Diprotodontia, Dasyuromorphia

Group2: Eulipotyphla, Perissodactyla, Pholidota

Group3: Macroscelidea, Pilosa, Proboscidea, Afrosoricida, Cingulata, Sirenia, Tubulidentata, Hyracoidea

Group 4: Lagomorpha, Rodentia, Scandentia Group 5: Primates, Dermoptera

Group 6: Artiodactyla

Group 7: Chiroptera

Group 8: Carnivora

We then added the most relevant, phylogenetically closest annotated *NEAT1* ortholog(s) to these groups and performed MSA. MSA was visualised using the online tool AlignmentViewer (https://alignmentviewer.org/). The coordinates of the genes’ start and stop sites (TATA-box and end of the triple helix) within the MSA were identified and used to build the final set of orthologs and their structural elements.The same procedure was applied for the *MALAT1* ortholog search, but sequences were divided into two groups: the aforementioned Group 1 and the remaining sequences.

Subsequently, we manually curated the results and removed orthologs with excessive assembly gaps or spurious sequences lacking the correct start or end. Coordinates, contig accessions, genome assembly versions, and other results and metadata can be found in Supplementary Table 1.

We examined the strand and genomic distance between *NEAT1* and *MALAT1*. Out of 428 organisms in which both genes were predicted, 92% had these genes located on the same contig. Since not all species possess complete chromosome-level assemblies, some genes were found on different contigs. In such cases, it is not possible to determine the true genomic positions of the genes. Among those located on the same contig, only two species had *NEAT1* and *MALAT1* coded on opposite strands. Assemblies of both these species, *Rousettus madagascariensis* and *Oryctolagus cuniculus*, do not belong to the GenBank reference set. After manual inspection, we found another reference assembly for *Oryctolagus cuniculus* (Suppl. Fig. 1B) and checked the strand and location of the predicted orthologs of *NEAT1* and *MALAT1*. Although these orthologs showed high sequence similarity to the reference assembly, the directionality of the genes was different: they were coded on the same strand, consistent with the majority of other orthologs. However, we cannot assess the impact of assembly quality on the opposing directionality observed in *Rousettus madagascariensis*, as no reference assembly is currently available.

### Comparison of naked mole-rat *Neat1* and *Malat1* to the results of Yamada *et al*

Our collection included a newer version of the naked mole-rat genome assembly than the one used in the publication by Yamada *et al*. (Yamada et al., 2022). We downloaded the assembly used in that study and performed a blastn search of the *Neat1* sequence identified in our study against this assembly. Our start coordinate for *Neat1* was 20,972,753, which is 204 bp downstream of the start coordinate reported by Yamada *et al*. (JH602080:20,972,549, with both genes coded on the minus strand). The start coordinate for *Malat1* was 20,907,466, which is 60 bp downstream of the coordinate reported by Yamada *et al*. (JH602080:20,907,406). We assume that the 3’-ends of the genes in the naked mole-rat are identical to those we identified, as Yamada *et al*. also defined them computationally based on the similarity of triple helix and tRNA-like motifs.

### Prediction of short isoform in *NEAT1* orthologs

We divided the orthologs into two similarity groups, with marsupials and Monotremata (Group 1) sequences placed separately. The remaining orthologs were subjected to MSA (MAFFT, default parameters). We identified the position in the MSA corresponding to the PAS of human *NEAT1Short* and searched for the predicted PAS in the vicinity of this position in the orthologs. Polyadenylation signals were predicted by searching for the canonical motif ‘AATAAA’. If a single signal was detected within 110 bp (in both directions) of the PAS position in human *NEAT1*, it was considered an active PAS for the *NEAT1Short* orthologs.

In Group 1, we searched for two PASs using the same logic, based on the predicted sites for *Monodelphis domestica* (Cornelis et al., 2016a). In this prediction, there were three orthologs where we could not identify a single alternative polyadenylation signal, and these were omitted from the analysis.

### Prediction of sequence elements in *NEAT1* and *MALAT1* orthologs

We used DFAM database (downloaded on April 2022, (Storer et al., 2021)) of transposable elements and searched for similarities using blastn algorithm and applying 80-80-80 rule (at least 80bp of alignment length with 80% of nucleotide identity over alignment covering at least 80% of the TE).

Short sequence repeats were predicted with MISA-web ((Beier et al., 2017), default parameters).

We retrieved 1kb of promoter sequence for each ortholog of *NEAT1* and *MALAT1* and predicted transcription factors binding sites using FIMO tool ((Grant et al., 2011), part of MEME package v5.0.5,(Bailey et al., 2015)) and JASPAR database (core part, version 2022, vertebrates, (Rauluseviciute et al., 2024)). Sites with p-value < 10^−4^ were considered. GO terms were downloaded on October 2021 ((Ashburner et al., 2000; The Gene Ontology Consortium et al., 2023)), each gene was associated with all connected to it terms.

G-quadruplexes were predicted using pqsfinder R package (v.2.2.0, (Hon et al., 2017)).

Kmers were counted using in house script.

Self-complementary regions were assessed from the BLASTn search against the ortholog itself, and only reverse complementary hits were counted.

### Average nucleotide identity calculation

ANI between two sequences was calculated by using all blastn hits longer than 100bp and following the formula:

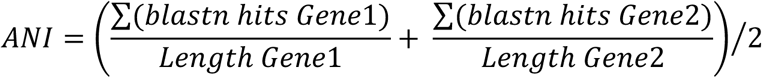

### Analysis of CDS and UTR regions of protein-coding gene orthologs in mammals

CDS and UTR regions of protein-coding gene orthologs in mammals were retrieved from GenBank (Clark et al., 2016) in September 2024 using the NCBI Datasets tool ((Na et al., 2024), command: datasets download gene symbol “$GENE” --ortholog mammals --include gene,cds,3p-utr,product-report; for genes with at least 200 orthologs from different genera). In cases where multiple transcripts were available, the longest single transcript per ortholog was selected. ANI, G-quadruplexes, and nucleotide usage were predicted in the same manner as for *NEAT1* and *MALAT1* orthologs; values were averaged across all orthologs per gene before being used in distribution plots. A total of 15,461 protein-coding genes were included in the CDS analysis, and 13,847 genes were used in the UTR analysis.

### Phylogenetic tree

The phylogenetic tree of Ns *et al*. (Ns et al., 2019) was used. Species for which *NEAT1* and *MALAT1* orthologs were identified but absent in the phylogenetic tree were associated with their closest relatives. A full list of these connections can be found in the ScriptName (GitHub link). Visualisation and graphical adjustments of the phylogenetic tree were made using the iTOL web server (Letunic and Bork, 2024). Tree parsing, pruning, and the retrieval of time information were performed using the ete3 Python package (v.3.1.2, (Huerta-Cepas et al., 2016)).

### Other used resources and software

RNAfold web-server ((Lorenz et al., 2011)) was used to predict and visualise folding of structural elements. LocRNA software (v. 2.0.0, http://rna.informatik.uni-freiburg.de, (Will et al., 2012)) was used to analyse coevolutionary patterns of tRNA-like structures. Sequence logos were generated using WebLogo web-server ((Crooks et al., 2004)). Taxonomic tree of NCBI (downloaded on April 2022, (Schoch et al., 2020)) was used to classify the studied genomes.

### Code and data availability

Most of analysis was performed with customs scripts, which were written in Python 3.7.0 and used the following packages: scipy (v.1.7.1, (Virtanen et al., 2020)), numpy (v. 1.18.5, (Harris et al., 2020)), pandas (v 1.1.5, https://zenodo.org/records/10957263), matplotlib (v.3.4.3, (Hunter, 2007)), seaborn (v. 0.11.2, (Waskom, 2021)) and Jupyter notebook (v.4.8.1, (Kluyver et al., 2016)). Code is available on GitHub (link). Orthologs sequences are deposited (link).

## Author contributions

K.A performed the research and prepared the manuscript, M.D. conceived the study and prepared the manuscript.

## Acknowledgements

We would like to thank the John Templeton Foundation for their generous support through Grant #62572.

## Declarations of interest

The authors declare no competing interests or financial conflicts related to this research.

---

**Supplementary Figure 1.**
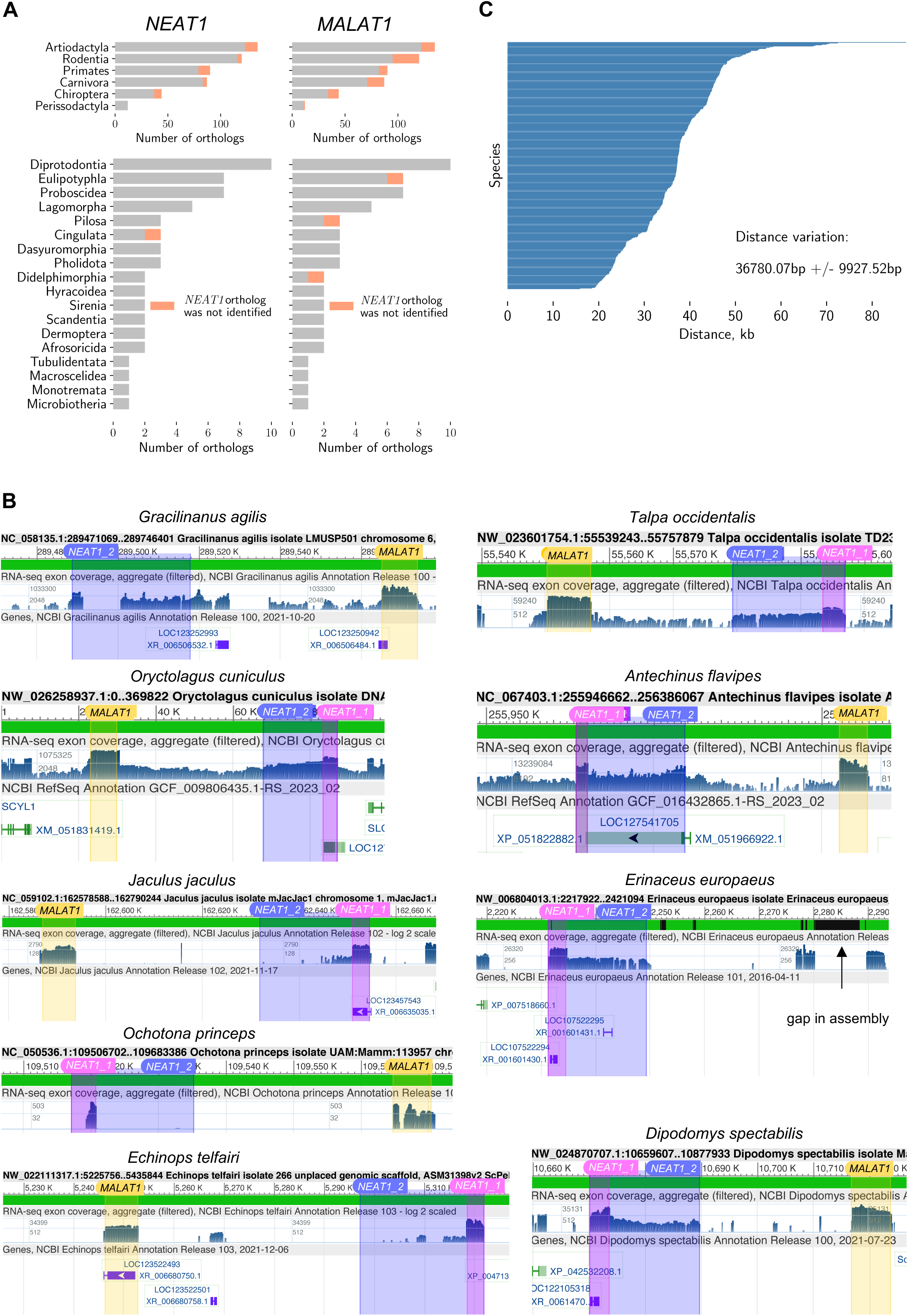
Identification of *NEAT1* and *MALAT1* orthologs in mammals. **A.** Number of identified orthologs in different mammalian orders. Species in which *NEAT1* or *MALAT1* orthologs were not found are highlighted in red. **B.** Transcription in regions with predicted *NEAT1* and *MALAT1* orthologs for species coding *NEAT1* archetypes. Predicted coordinates were overlaid on mapped transcriptomic read profiles in the Genome Browser (http://genome.ucsc.edu). In the ‘Genes’ section of the Genome Browser, the automatically predicted genes identified in the region are shown. **C.** Distribution of the genomic distance between *NEAT1* and *MALAT1* in studied mammals.

**Supplementary Figure 2.**
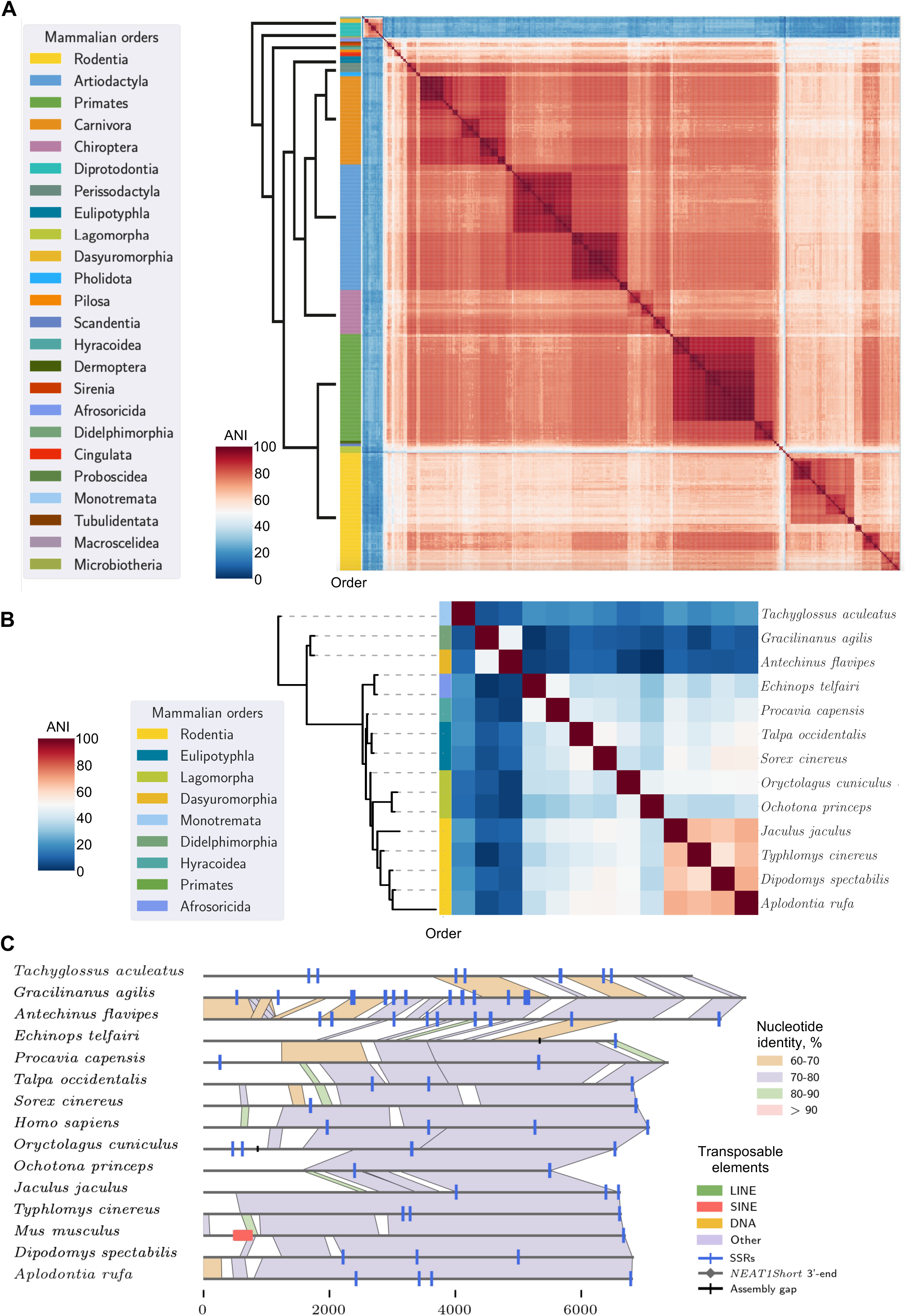
Primary sequence diversity of *MALAT1* orthologs in all mammals and the subset of archetypes. **A.** Heatmap of ortholog similarity in pairwise comparisons (all-to-all). Orthologs are arranged along a phylogenetic tree, and the colour bar on the left corresponds to the mammalian orders of individual orthologs; colours are explained in the legend. For visual clarity, the phylogenetic tree was simplified, and information about phylogenetically estimated time was omitted. Red clusters represent groups of highly similar orthologs, while blue areas indicate lower similarity between them. **B.** Heatmap of primary sequence similarity among *MALAT1* orthologs coding *NEAT1* archetypes. Orthologs are arranged along a phylogenetic tree, and the colour bar on the left corresponds to the mammalian orders of individual orthologs; colours are explained in the legend. The phylogenetic tree is time-scaled. *MALAT1* orthologs in the subset are also among the most diverged. **C.** Graphical representation of pairwise alignment between *MALAT1* archetypes. Patches of similarity longer than 100bp are depicted.

**Supplementary Figure 3.**
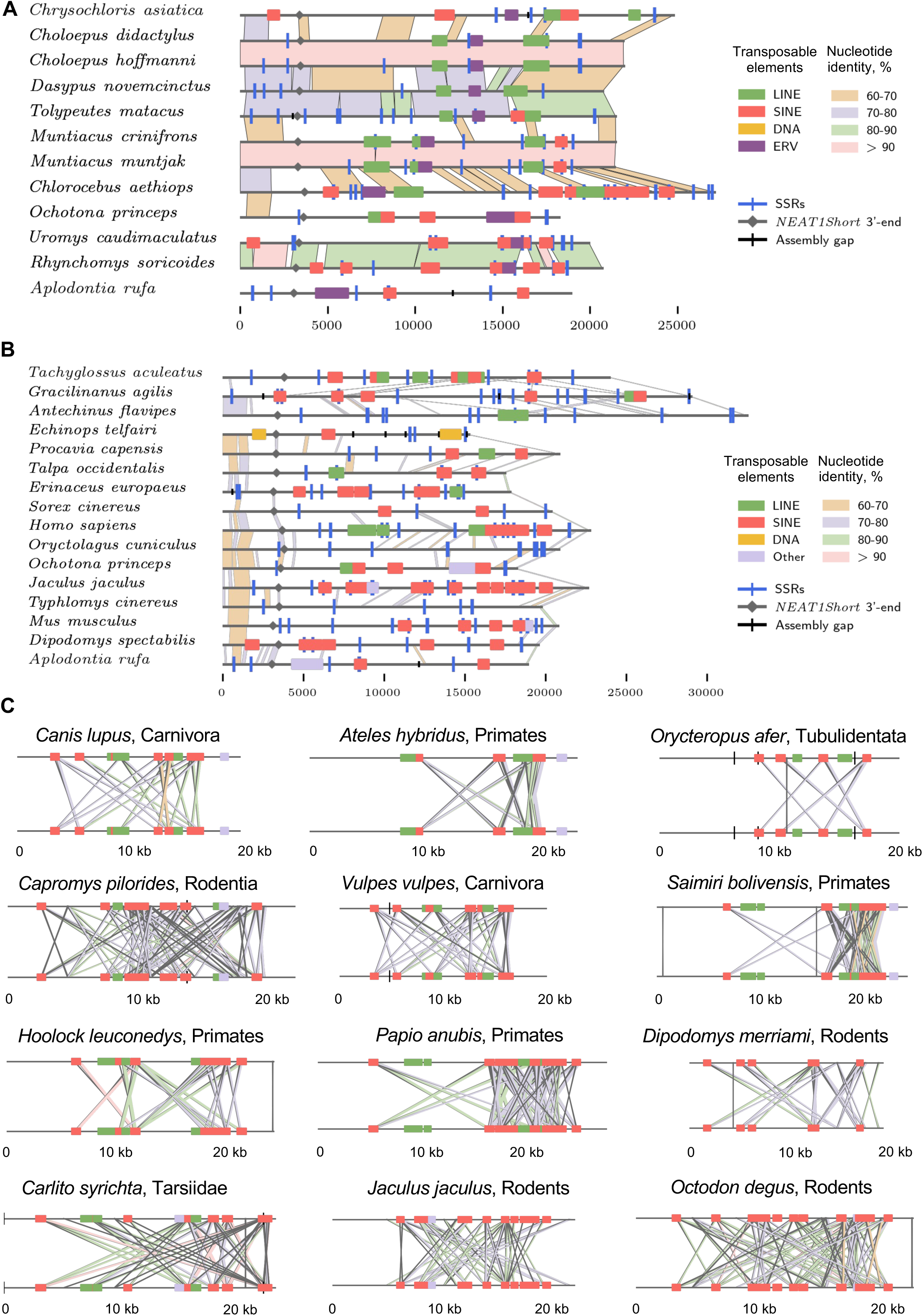
Transposable elements in *NEAT1* orthologs. **A.** Pairwise alignment of *NEAT1* orthologs carrying ERV-type TEs. Genomes are aligned along the phylogenetic tree. Only patches of similarity longer than 500bp are depicted. **B.** Pairwise alignment of *NEAT1* archetypes, arranged along the phylogenetic tree. Patches of similarity longer than 100bp are depicted. **C.** Graphical representation of the distribution of self-complementary regions in selected *NEAT1* orthologs. Each ortholog is aligned to itself, and reverse complementary regions are connected with lines, forming a visual ‘cross’ shape. Orthologs were selected to demonstrate the full diversity of possible distributions of self-complementary regions. Cases lacking such regions were not included.

**Supplementary Figure 4.**
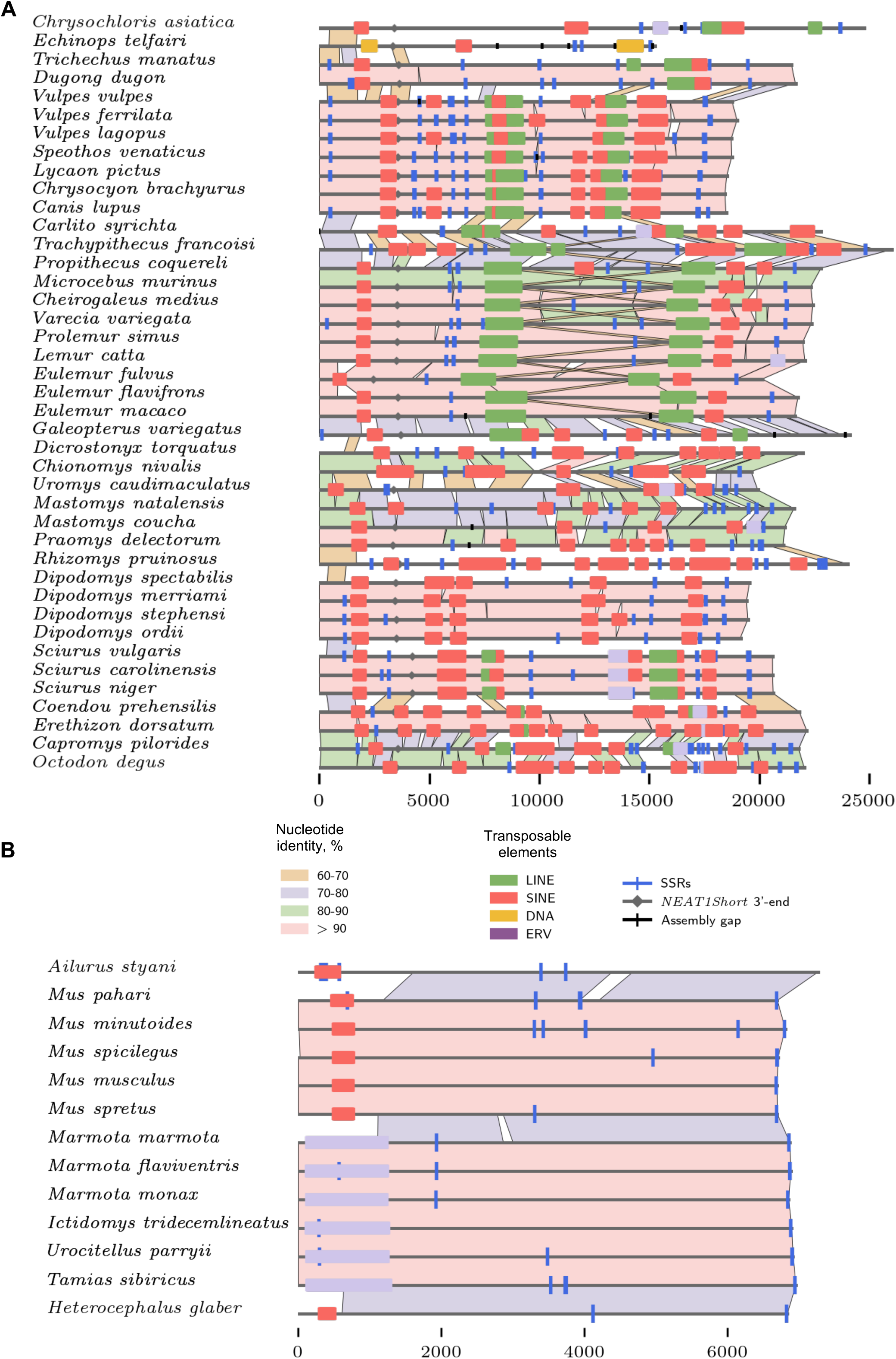
Transposable elements in *NEAT1Short* and *MALAT1* orthologs. **A.** Pairwise alignment of *NEAT1* orthologs carrying TEs in the *NEAT1Short* isoform. Species are ordered according to the phylogenetic tree. In the figure, only one *NEAT1* ortholog of *Canis lupus* from the eight different assemblies in our dataset is presented. The alignment of all eight orthologs of *Canis lupus* can be found in Suppl. Fig. 9. **B.** *MALAT1* orthologs carrying TEs. Species are ordered according to the phylogenetic tree.

**Supplementary Figure 5.**
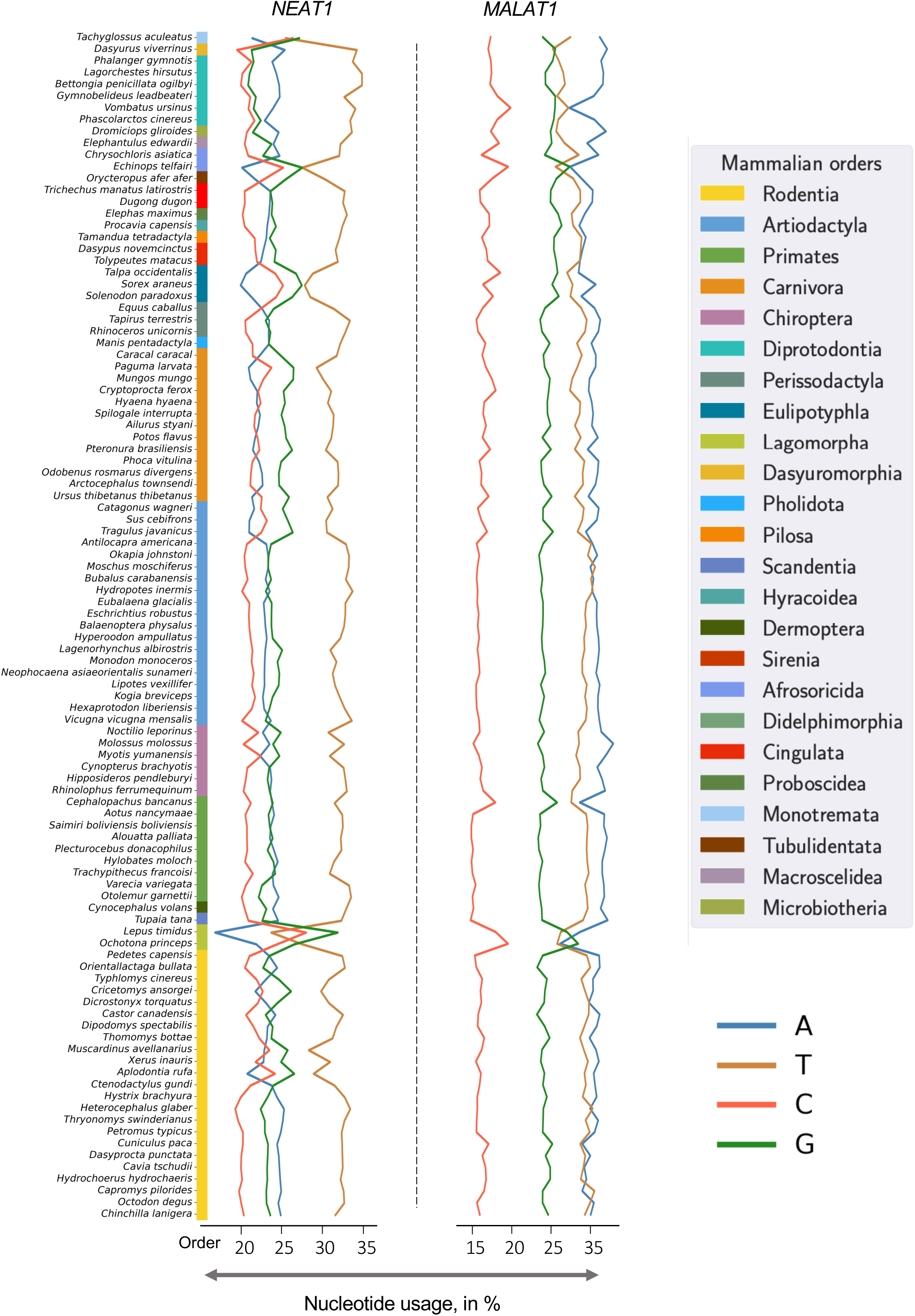
Nucleotide usage of *NEAT1* and *MALAT1* orthologs across mammalian orders. One species per family is depicted. The colour bar corresponds to mammalian orders, arranged along the phylogenetic tree.

**Supplementary Figure 6.**
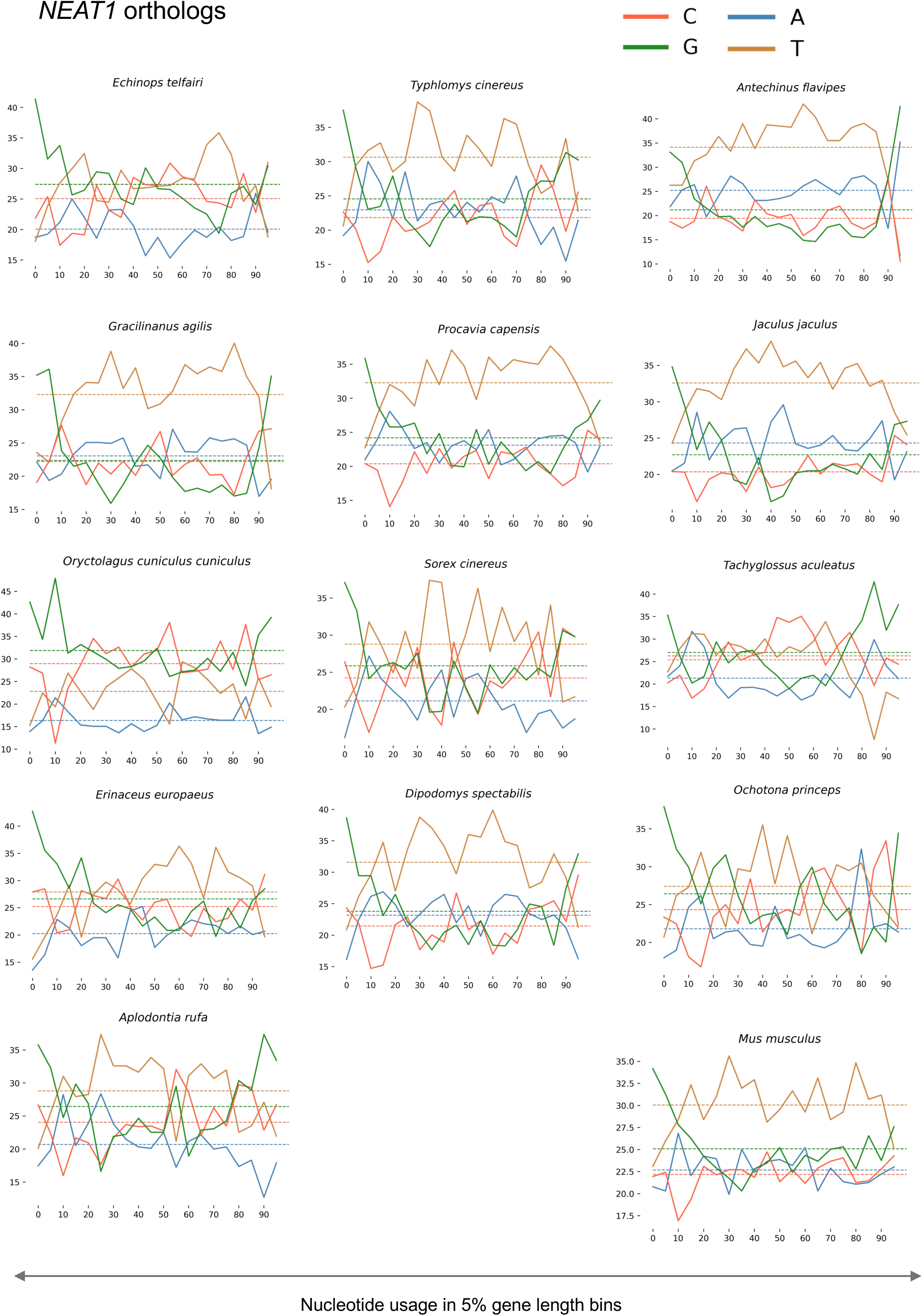
Plot of nucleotide usage along the sequence for *NEAT1* archetypes. The length of the ortholog was binned into 5% segments, and the nucleotide usage of each bin was estimated. The average nucleotide usage is depicted with dashed lines.

**Supplementary Figure 7.**
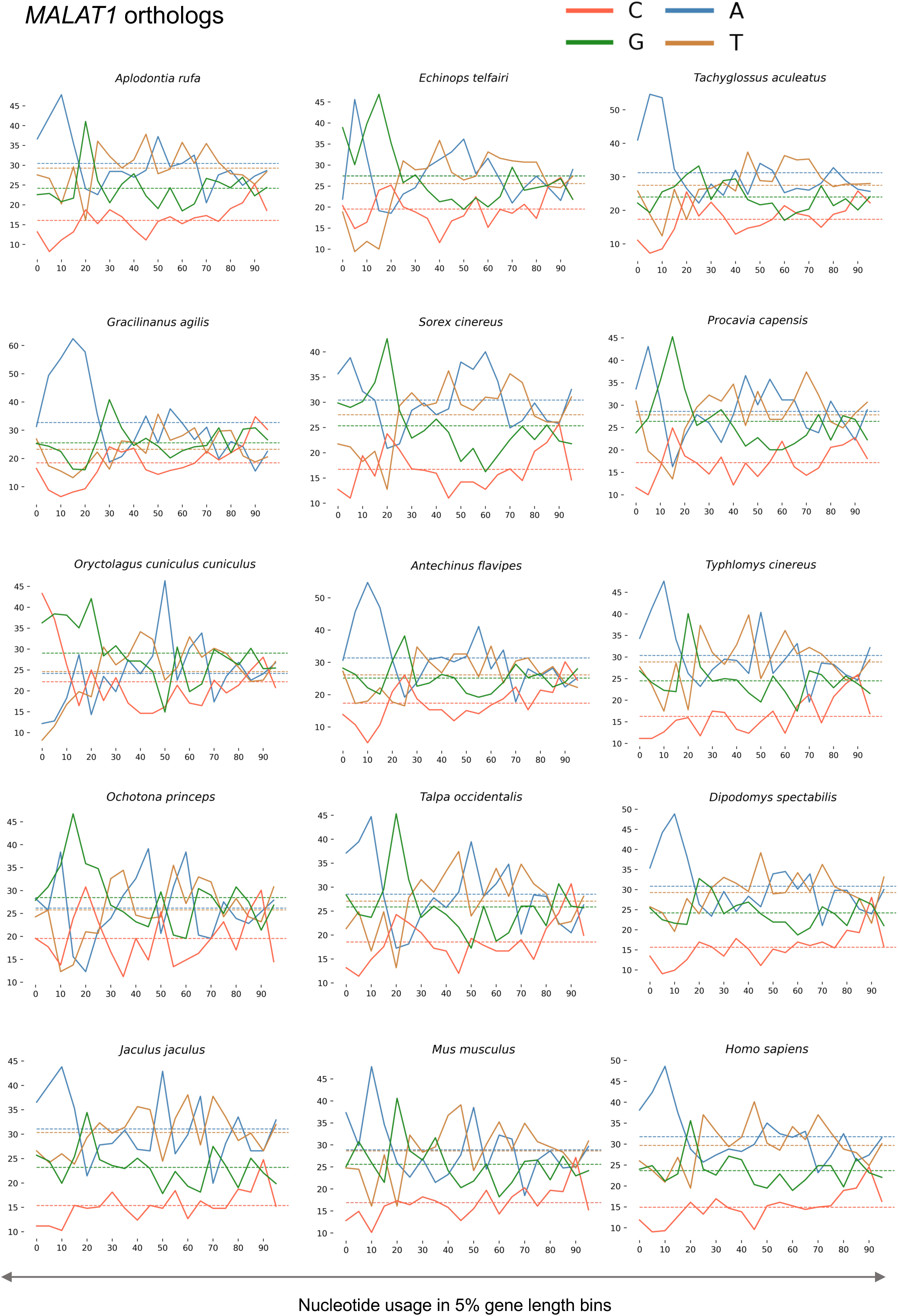
Plot of nucleotide usage along the sequence for *MALAT1* archetypes. The length of the ortholog was binned into 5% segments, and the nucleotide usage of each bin was estimated. The average nucleotide usage is depicted with dashed lines.

**Supplementary Figure 8.**
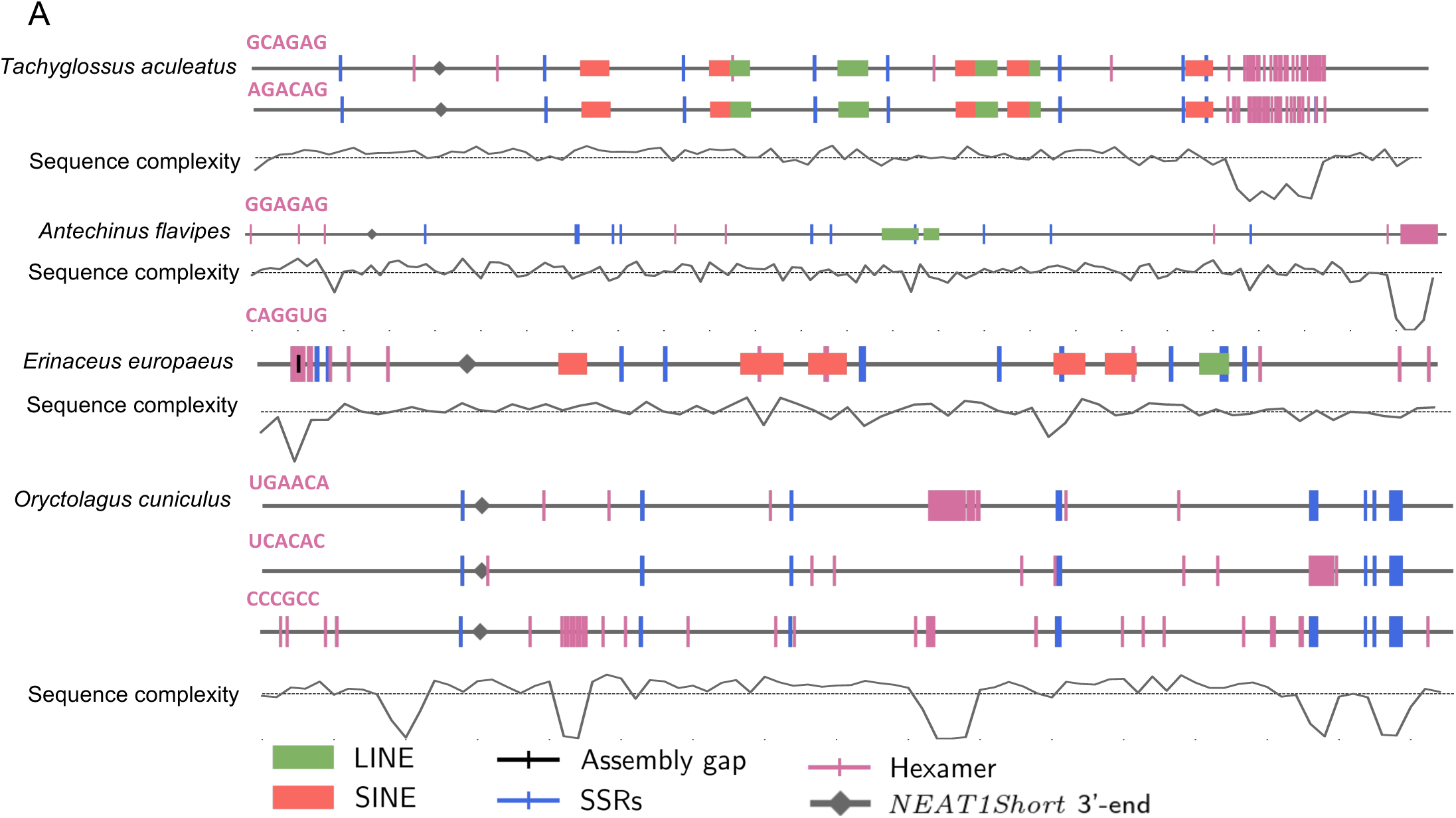
Distribution of the most frequent hexamers specific to an individual *NEAT1* ortholog from the archetypes subset. The depicted hexamers are listed on the left side of the ortholog map. Each vertical line, in the colours described in the legend, marks the position of the identified element. Ortholog maps are paired with the sequence complexity plot, highlighting drops in complexity caused by clustering of hexamer repeats.

**Supplementary Figure 9.**
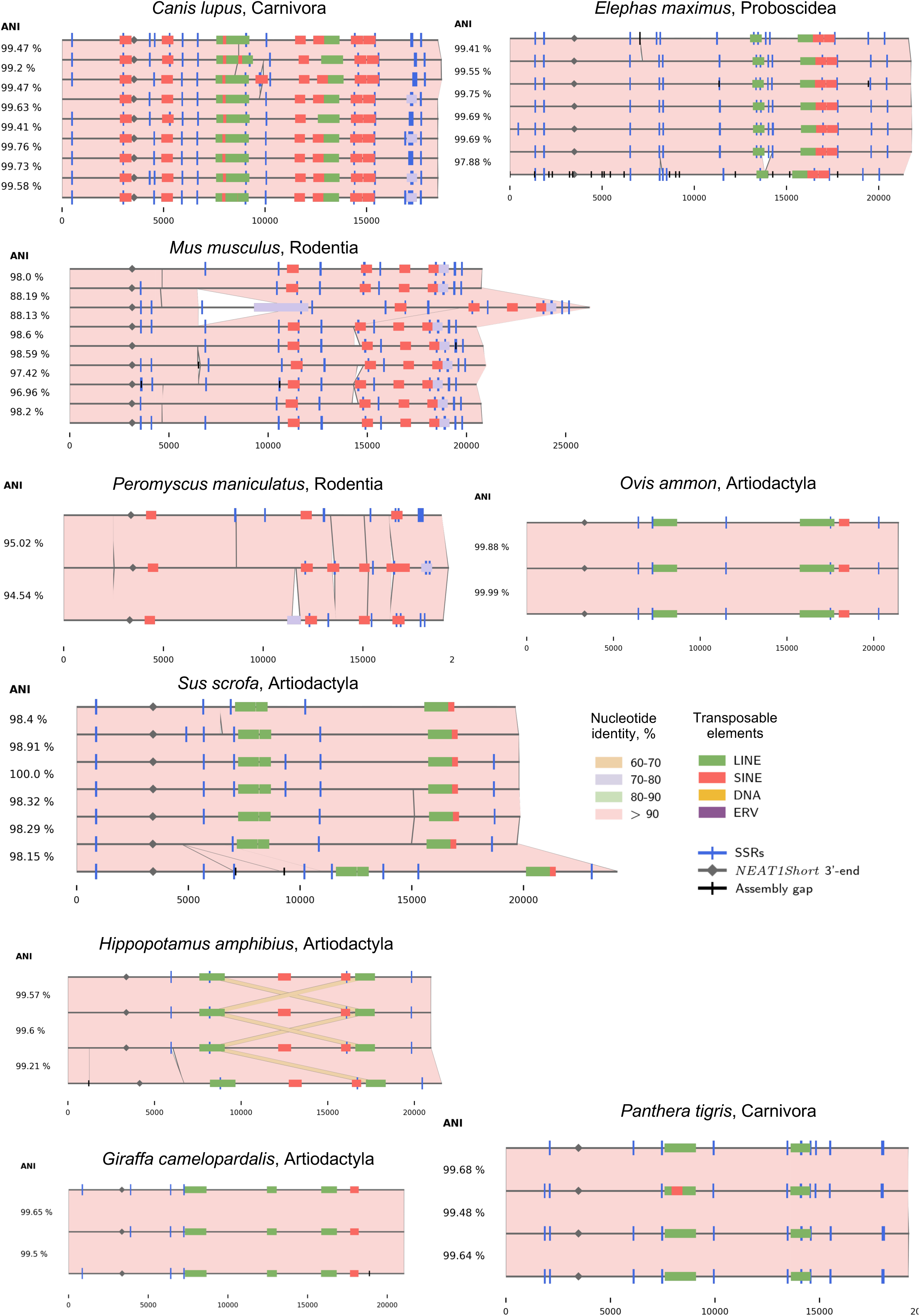
Pairwise alignments of *NEAT1* orthologs from different assemblies of the same species. On the left side, the ANI value of the pairwise comparison is indicated.

**Supplementary Figure 10.**
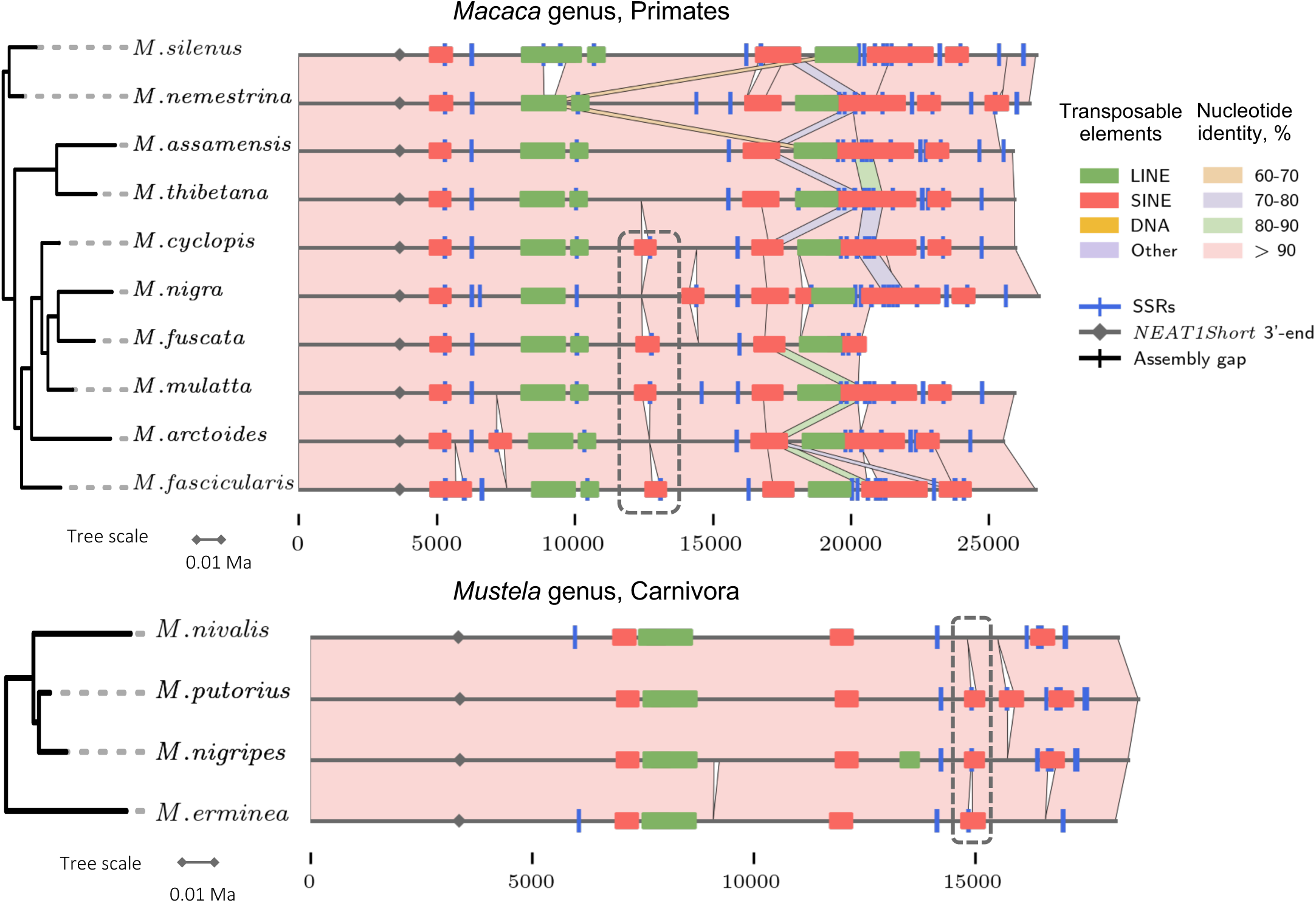
Instances of detected excised SINE elements. Pairwise alignment of *Macaca* and *Mustela* genera. Regions with deleted SINE elements are highlighted by a dashed frame.

## Notes

### Competing Interest Statement

The authors have declared no competing interest.

